# Structural and Evolutionary Constraints of Organophosphate Resistance in Dipteran Carboxylesterases

**DOI:** 10.1101/2025.07.07.663592

**Authors:** Rebecca L. Frkic, Alex Giang, Jian-Wei Liu, Mojtaba Esmaeily, Paul D. Carr, Nicholas J. Fraser, Davis Hopkins, John G. Oakeshott, Philip Batterham, Peter D. Mabbitt, Colin J. Jackson

## Abstract

Enzymatic detoxification of organophosphate (OP) insecticides can confer resistance in some insects, yet the precise molecular basis of this trait, and how it has evolved, remains poorly understood. In certain dipteran species, a G→D mutation in the oxyanion hole of α-carboxylesterases (CBEs) enhances OP hydrolysis, yet this adaptation is not widespread despite the presence of orthologous CBEs in other insect species that are also exposed to OPs. The extent, and molecular basis, of evolutionary contingency and epistasis in this catalytic OP resistance has not been explored, and how further mutations might optimize OP detoxification in the future is not clear. Here, we systematically compare OP hydrolysis and analyse structures of CBE orthologs across several dipteran species, revealing that the success of the G137D mutation is sequence context-dependent. We employed laboratory-directed evolution to enhance OP turnover over 1000-fold *vs.* the wild-type enzyme and tested these variants in transgenic *Drosophila melanogaster*, demonstrating that improved catalytic rates do not directly translate to increased resistance. By highlighting the trade-off between organophosphate affinity and turnover, this work further clarifies the complex evolutionary trajectories determining why a particular resistance mechanism may evolve in some species but not others.

**Significance:** This study reveals the intricate evolutionary path to insecticide resistance in insects, highlighting why a potent resistance mutation is effective in some species but not others. We show that the mutation’s success is contingent on the enzyme’s pre-existing structural features, highlighting the strong intramolecular epistasis. Using laboratory evolution, we enhanced the enzyme’s detoxification activity over 1000-fold, yet discovered this did not translate to increased resistance in transgenic flies. This surprising result demonstrates that effective real-world resistance requires a delicate balance between an enzyme’s ability to bind an insecticide (affinity) and its speed at breaking it down (turnover), providing crucial insights into the constraints governing molecular adaptation.

## Introduction

Organophosphate (OP) insecticides have been deployed globally to control insect populations by targeting acetylcholinesterase (AChE), a critical enzyme in nerve signal transduction (1). However, catalytic (enzyme-based) resistance in insects emerged rapidly, which has led to interest in the genetic and biochemical adaptations underlying detoxification (2–4). The Australian sheep blowfly, *Lucilia cuprina*, is one of the best studied instances of the evolution of catalytic OP resistance, where a single amino acid substitution (Gly137Asp) in the oxyanion hole of the active site in the α-carboxylesterase 7 (*α*E7) enzyme confers resistance to OPs such as diazinon (5) (**Supplementary Figure 1**). The Gly137Asp (G137D) mutation in *α*E7 is an example of neofunctionalization, with the enzyme losing most of its native carboxylesterase activity and becoming an OP-hydrolase, and analogous single amino acid substitutions repurposing orthologous carboxylesterases (CBEs) have been observed in other dipteran insects (6–8). While the G137D class of mutations can compromise native CBE activity and reduce fitness, their selective advantage under intense OP pressure has enabled the mutation to remain in populations around the world with the subsequent evolution of epistatic modifier mutations and gene duplication to compensate for the loss of the native activity (9–11). Parallel adaptations, including Trp251Leu in *L. cuprina* or Trp251Ser substitutions in *Cochliomyia hominivorax* and analogous mutations in *Musca domestica* have also been observed (6, 12–14), although the tradeoff between the new and original activity in these examples is much lower (9). The precise structural and epistatic constraints shaping *α*E7 evolution under OP selection pressure remain unresolved, highlighting the need to dissect the interplay between sequence context, catalytic efficiency, and ecological fitness.

From a mechanistic standpoint, OP insecticides exert their toxicity by phosphorylating the catalytic serine in serine hydrolases, including the primary target, acetylcholinesterase, and many CBEs (15). This modification inactivates these enzymes, leading to interminable nerve signal transduction and death (16). However, certain CBEs can sequester and, in some instances, hydrolyse OP compounds, mitigating their toxic effects (17–21). In *α*E7 from *L. cuprina*, the G137D substitution introduces a general base into the active site, allowing for catalytic turnover of OP substrates, rather than mere sequestration (5, 22, 23). Structural analyses of *α*E7 have shown that Asp137 can facilitate dephosphorylation of the serine:OP intermediate, a necessary step for restoring the active enzyme (22, 23). However, this step requires precise alignment of adjacent residues and dynamic rearrangements within the active site. Structural and dynamical analysis has revealed the importance of ancillary residues such as Met308 and Phe309, which help orient Asp137 into a catalytically productive conformation (20, 22). Although it increases catalytic turnover, the G137D mutation introduces steric and electrostatic clashes with OP substrates, and often increases the Michaelis constant (*K*_m_) and reduces affinity (22, 24, 25). This balance between substrate affinity and catalytic turnover underpins many of the complexities in the structural and biochemical basis of OP resistance.

CBE-mediated OP resistance in insects is a useful real-world, eukaryotic system to study evolutionary contingency and epistasis in protein evolution. In evolution, permissive or “neutral” substitutions can accumulate over time, unpredictably affecting the accumulation of future adaptive mutations (26–29). However, due to the context dependence of mutations, or epistasis, mutations that might work in one species’ enzyme scaffold may not be viable in another’s (30, 31), i.e. the effects of mutations are contingent on the evolutionary histories of the CBEs in various species (32). These concepts have been well studied in laboratory systems in prokaryotes, e.g., the evolution of antibiotic resistance in bacterial β-lactamases (27, 33). However, the observation that the G137D mutation has occurred in a fraction of insect species, yet has not become fixed in many more in populations across the globe (5–7), suggests that epistasis and evolutionary contingency might be important factors in its evolution. In other words, *α*E7 from certain species may be “primed” to accommodate Asp137, whereas other species appear to lack the pre-existing conditions necessary for Asp137-based resistance.

Despite a growing understanding of CBE-mediated OP resistance in insects, key questions remain unanswered. First, it is not clear why only a subset of dipteran species efficiently exploit the G137D (*L. cuprina α*E7 numbering henceforth) pathway while others do not, even though they share similar ecological niches and face analogous selection pressures. Second, the *in vivo* significance of enhanced OP-hydrolysis rates is still under debate: will catalytic improvements translate into field-level resistance if other aspects of function, such as affinity, are compromised. Third, what does the future hold in store for this resistance mechanism? Now that the enzyme has been neofunctionalized into an OP hydrolase, should we expect its activity to continue to improve? In this work, we address these questions by (1) systematically comparing *α*E7 orthologs from diverse dipteran species to discern the structural motifs, binding networks, and conformational dynamics that enhance or restrict the evolution of OP hydrolysis; (2) employing laboratory-directed evolution on *L. cuprina α*E7 to identify additional substitutions that might increase OP hydrolase activity beyond what is currently observed in the wild; and (3) testing the *in vivo* relevance of newly evolved variants generating transgenic *Drosophila melanogaster* strains and investigating their resistance to OPs. Through these approaches, we reveal (i) how subtle sequence features and conformational states shape the evolutionary trajectories of OP resistance, (ii) provide new insight into why some species evolve potent detoxification strategies while others remain confined to less efficient resistance mechanisms, and (iii) demonstrate that resistance is more complex than *in vitro* enzyme kinetics with the relative importance of sequestration and turnover being balanced.

## Results and Discussion

### The molecular and biochemical effects of the G137D mutation are dependent on genetic context

We began by surveying the steady-state hydrolytic activities of α-carboxylesterase 7 (*α*E7) orthologs from nine dipteran species with varying degrees of similarity to *L. cuprina α*E7 (89-38% amino acid identity; **Table 1, Supplementary Figure 2**). These orthologs were selected to encompass a broad range of insect families that have historically encountered organophosphate (OP) insecticides, from blowflies to mosquitoes (**Table 1**). Using the model carboxylester substrate 4-nitrophenyl butyrate (4-NPB), we observed that introducing the G137D mutation consistently reduced native esterase turnover (*k*_cat_) across most species, reflecting an inherent strong-positive trade-off between native function and OP detoxification and that the G137D mutation in the oxyanion hole of *α*E7 enzymes is deleterious to the native activity (**Table 2**). This stands in contrast to other examples of enzyme neofunctionalization, where weak negative tradeoffs, i.e. a promiscuous activity can increase without large losses of the native activity, are often seen (34, 35).

**Table 1:**
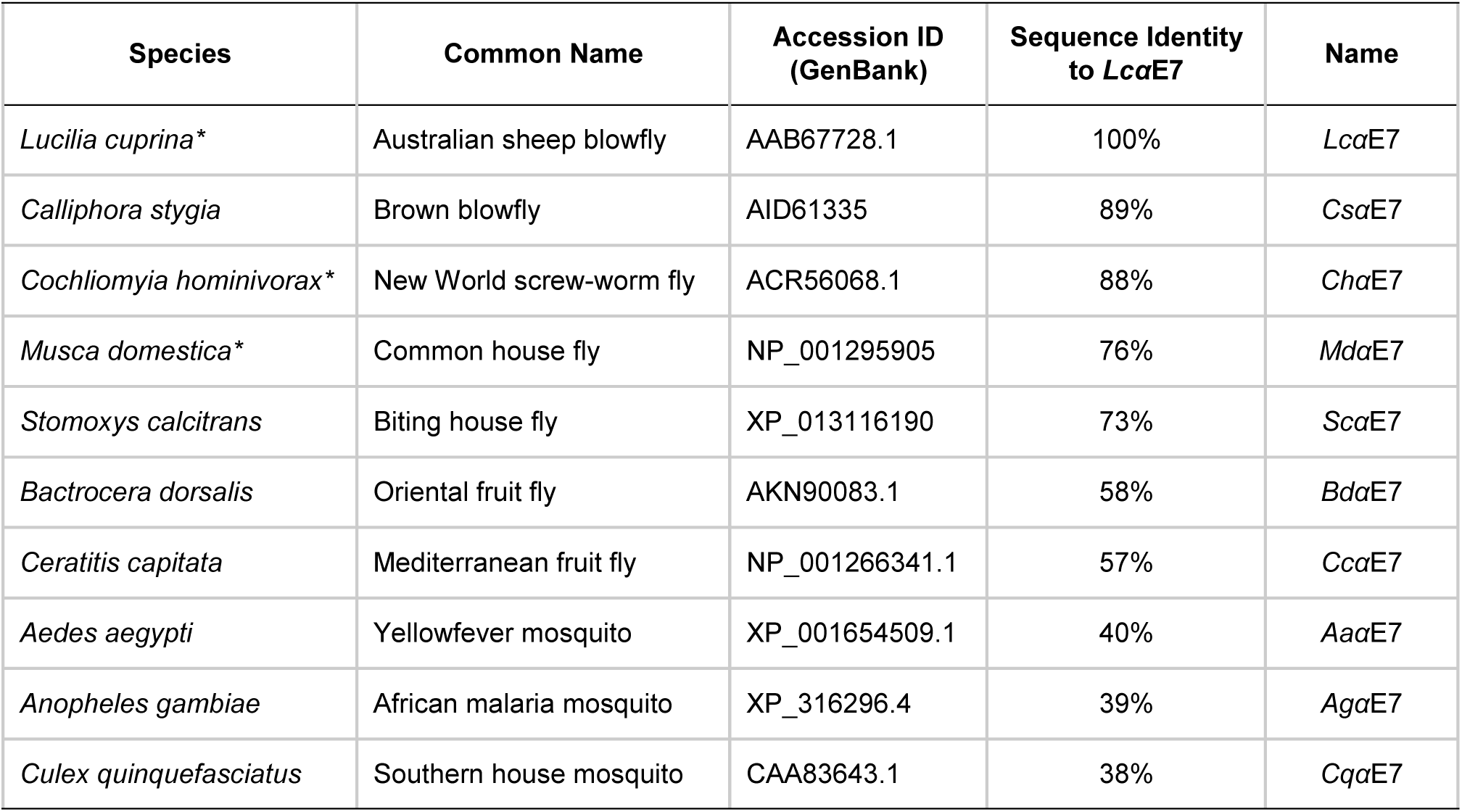
Dipteran *α*E7 orthologs investigated in this study. Summary of the *α*E7 orthologs from various dipteran species studied in this work. * Indicates species that have acquired the G137D mutation in nature.

| Species | Common Name | Accession ID (GenBank) | Sequence Identity to *LcaE7* | Name |
| --- | --- | --- | --- | --- |
| *Lucilia cuprina* * | Australian sheep blowfly | AAB67728.1 | 100% | *LcaE7* |
| *Calliphora stygia* | Brown blowfly | AID61335 | 89% | *CsaE7* |
| *Cochliomyia hominivorax* * | New World screw-worm fly | ACR56068.1 | 88% | *ChaE7* |
| *Musca domestica* * | Common house fly | NP_001295905 | 76% | *MdaE7* |
| *Stomoxys calcitrans* | Biting house fly | XP_013116190 | 73% | *ScaE7* |
| *Bactrocera dorsalis* | Oriental fruit fly | AKN90083.1 | 58% | *BdaE7* |
| *Ceratitis capitata* | Mediterranean fruit fly | NP_001266341.1 | 57% | *CcaE7* |
| *Aedes aegypti* | Yellowfever mosquito | XP_001654509.1 | 40% | *AaaE7* |
| *Anopheles gambiae* | African malaria mosquito | XP_316296.4 | 39% | *AgαE7* |
| *Culex quinquefasciatus* | Southern house mosquito | CAA83643.1 | 38% | *CqaE7* |

**Table 2.**
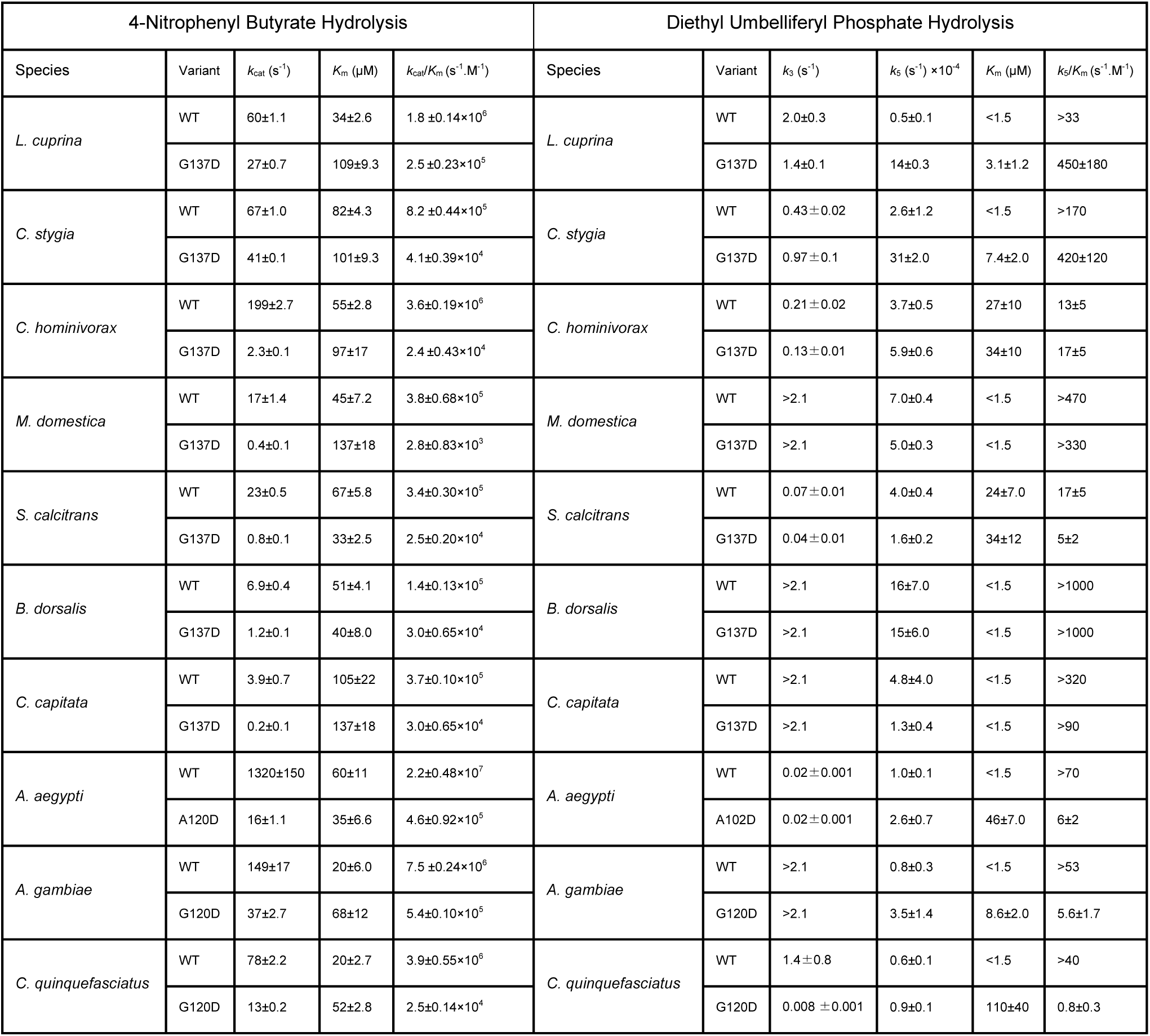
Michaelis-Menten kinetic parameters for hydrolysis of 4-nitrophenyl butyrate (carboxylester), and diethyl umbelliferyl phosphate (organophosphate). Values are mean ± standard error (n=3).

| 4-Nitrophenyl Butyrate Hydrolysis |  |  |  |  | Diethyl Umbelliferyl Phosphate Hydrolysis |  |  |  |  |  |
| --- | --- | --- | --- | --- | --- | --- | --- | --- | --- | --- |
| Species | Variant | $k_{cat}$ (s<sup>-1</sup>) | $K_m$ (μM) | $k_{cat}/K_m$ (s<sup>-1</sup>.M<sup>-1</sup>) | Species | Variant | $k_3$ (s<sup>-1</sup>) | $k_5$ (s<sup>-1</sup>) ×10<sup>-4</sup> | $K_m$ (μM) | $k_5/K_m$ (s<sup>-1</sup>.M<sup>-1</sup>) |
| *L. cuprina* | WT | 60±1.1 | 34±2.6 | 1.8 ±0.14×10<sup>6</sup> | *L. cuprina* | WT | 2.0±0.3 | 0.5±0.1 | <1.5 | >33 |
|  | G137D | 27±0.7 | 109±9.3 | 2.5 ±0.23×10<sup>5</sup> |  | G137D | 1.4±0.1 | 14±0.3 | 3.1±1.2 | 450±180 |
| *C. stygia* | WT | 67±1.0 | 82±4.3 | 8.2 ±0.44×10<sup>5</sup> | *C. stygia* | WT | 0.43 ±0.02 | 2.6±1.2 | <1.5 | >170 |
|  | G137D | 41±0.1 | 101±9.3 | 4.1±0.39×10<sup>4</sup> |  | G137D | 0.97 ±0.1 | 31±2.0 | 7.4±2.0 | 420±120 |
| *C. hominivorax* | WT | 199±2.7 | 55±2.8 | 3.6±0.19×10<sup>6</sup> | *C. hominivorax* | WT | 0.21 ±0.02 | 3.7±0.5 | 27±10 | 13±5 |
|  | G137D | 2.3±0.1 | 97±17 | 2.4 ±0.43×10<sup>4</sup> |  | G137D | 0.13 ±0.01 | 5.9±0.6 | 34±10 | 17±5 |
| *M. domestica* | WT | 17±1.4 | 45±7.2 | 3.8±0.68×10<sup>5</sup> | *M. domestica* | WT | >2.1 | 7.0±0.4 | <1.5 | >470 |
|  | G137D | 0.4±0.1 | 137±18 | 2.8±0.83×10<sup>3</sup> |  | G137D | >2.1 | 5.0±0.3 | <1.5 | >330 |
| *S. calcitrans* | WT | 23±0.5 | 67±5.8 | 3.4±0.30×10<sup>5</sup> | *S. calcitrans* | WT | 0.07 ±0.01 | 4.0±0.4 | 24±7.0 | 17±5 |
|  | G137D | 0.8±0.1 | 33±2.5 | 2.5±0.20×10<sup>4</sup> |  | G137D | 0.04 ±0.01 | 1.6±0.2 | 34±12 | 5±2 |
| *B. dorsalis* | WT | 6.9±0.4 | 51±4.1 | 1.4±0.13×10<sup>5</sup> | *B. dorsalis* | WT | >2.1 | 16±7.0 | <1.5 | >1000 |
|  | G137D | 1.2±0.1 | 40±8.0 | 3.0±0.65×10<sup>4</sup> |  | G137D | >2.1 | 15±6.0 | <1.5 | >1000 |
| *C. capitata* | WT | 3.9±0.7 | 105±22 | 3.7±0.10×10<sup>5</sup> | *C. capitata* | WT | >2.1 | 4.8±4.0 | <1.5 | >320 |
|  | G137D | 0.2±0.1 | 137±18 | 3.0±0.65×10<sup>4</sup> |  | G137D | >2.1 | 1.3±0.4 | <1.5 | >90 |
| *A. aegypti* | WT | 1320±150 | 60±11 | 2.2±0.48×10<sup>7</sup> | *A. aegypti* | WT | 0.02 ±0.001 | 1.0±0.1 | <1.5 | >70 |
|  | A120D | 16±1.1 | 35±6.6 | 4.6±0.92×10<sup>5</sup> |  | A102D | 0.02 ±0.001 | 2.6±0.7 | 46±7.0 | 6±2 |
| *A. gambiae* | WT | 149±17 | 20±6.0 | 7.5 ±0.24×10<sup>6</sup> | *A. gambiae* | WT | >2.1 | 0.8±0.3 | <1.5 | >53 |
|  | G120D | 37±2.7 | 68±12 | 5.4±0.10×10<sup>5</sup> |  | G120D | >2.1 | 3.5±1.4 | 8.6±2.0 | 5.6±1.7 |
| *C. quinquefasciatus* | WT | 78±2.2 | 20±2.7 | 3.9±0.55×10<sup>6</sup> | *C. quinquefasciatus* | WT | 1.4 ±0.8 | 0.6±0.1 | <1.5 | >40 |
|  | G120D | 13±0.2 | 52±2.8 | 2.5±0.14×10<sup>4</sup> |  | G120D | 0.008 ±0.001 | 0.9±0.1 | 110±40 | 0.8±0.3 |

To gain insight into the effects of the G137D mutation on the OP hydrolysis activity and the mechanistic underpinnings of these differences, we employed stopped-flow fluorescence kinetics, monitoring the rapid phosphorylation and subsequent dephosphorylation of the active-site serine (**Table 2**). By tracking the release of the fluorescent product, we could distinguish between the fast “burst” phase (*k*_3_), corresponding to phosphorylation, and the rate-limiting dephosphorylation step (*k*_5_) that determines catalytic turnover (**Figure 1**). For example, in the wild-type *Lcα*E7, we observe a rapid burst phase, in which OP binds at the active site and is hydrolyzed, forming what is essentially an irreversibly phosphorylated intermediate. Eventually every enzyme molecule is phosphorylated and the turnover rate is limited by the slower dephosphorylation step (*k*_5_). In contrast, in *Lcα*E7 G137D, we still observe a rapid burst phase (*k*_3_), but also detect a significant increase in *k*_5_ (0.5 to 14 s^-1^), indicating hydrolysis of the phospho-serine intermediate, regeneration of the active site and multiple turnovers. We also observe a large increase in *k*_5_ in the closely related (89% amino acid identity) *Csα*E7, but most other G137D orthologs showed only modest improvements in *k*_5_, or little change compared to wild-type. Notably, the distantly related *Cqα*E7 (38% amino acid identity), exhibited a marginal increase in *k*_5_, but a significant decrease in *k*_3_ and a large increase in *K*_m_ that suggests the mutation affects OP binding. These findings confirm that G137D imparts highly context-dependent benefits to OP-hydrolysis, demonstrating strong evolutionary contingency and emphasizing how sequence background and structural environment can govern a mutation’s selective advantage.

**Figure 1:**
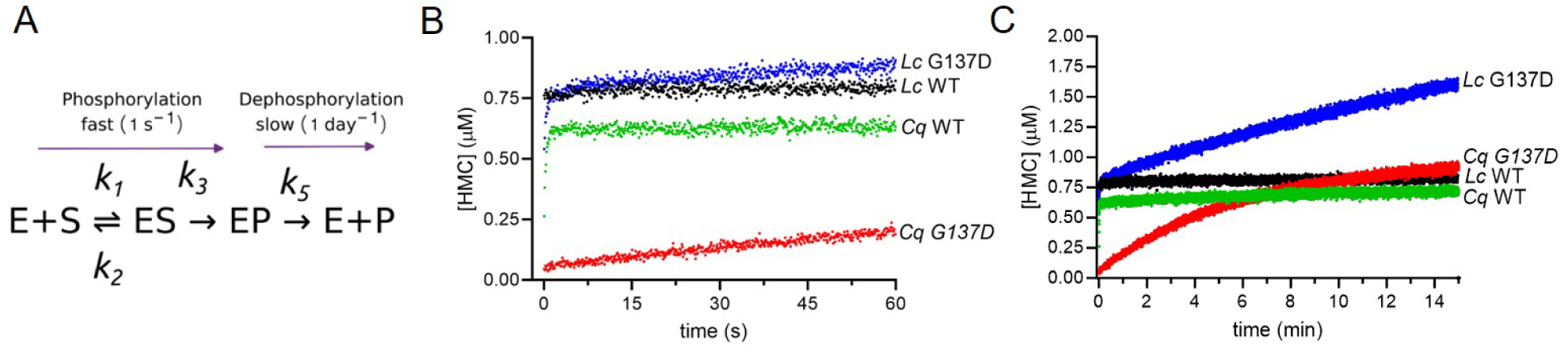
**A)** Kinetic scheme for OP-hydrolysis by *α*E7. **B)** Representative stopped-flow kinetic data for OP-hydrolysis by WT and G137D *L. cuprina* and *C. quinquefasciatus α*E7 over 1 minute, or **C)** 15 minutes. The concentration of product HMC (7-hydroxy-4-methyl coumarin) is indicated.

To obtain structural insight into the context-dependence of the G137D substitution, we solved the X-ray crystal structures of the mosquito *Anopheles gambii Agα*E7 and the New World screw-worm *Cochliomylia hominivorax Chα*E7 wild type enzymes (**Supplementary Table 1**), and analyzed them in combination with previously solved crystal structures of *Lucilia cuprina Lcα*E7 wild type (PDB 4FNG)(23), *Lcα*E7 G137D (PDB 5C8V)(22) and *Culex quinquefasciatus* Cq*α*E7 wild type (PDB 5W1U)(17) (**Figure 2A**). Thus, we have empirical structures of *Lcα*E7 and orthologs ranging from 88% amino acid identity (*Chα*E7) and moderate acceptance of the G137D mutation, to 39% (*Agα*E7) and 38% (*Cqα*E7) and poor acceptance of the G137D mutation (**Tables 1 & 2**). Structural alignment enabled comparison of active site residues; an overview of amino acid identities within the active site can be found in **Supplementary Table 2**. In *Lcα*E7, the D137 side chain participates in the hydrolytic dephosphorylation by stabilizing an attacking water molecule, but this favorable arrangement relies on a network of ancillary residues that orient Asp137 and the catalytic water (22). Notably, residues near the active site, such as M308 and F309, are able to shift their conformations to accommodate the bulkier aspartate side chain, but equally can constrain it to a catalytically productive orientation, thereby enhancing turnover (*k*_5_) (**Figure 2B**). Concurrently, Tyr457 is ideally positioned to form a water-mediated hydrogen-bond network with Asp137 (**Figure 2C**). In contrast, orthologs with different residues at positions corresponding to 308 and 309, such as F308 and S309 in *Cqα*E7, result in significantly different active site that does not support Asp137 in its catalytically productive rotamer, leading to an inability to efficiently orient the attacking water molecule for dephosphorylation. Similarly, certain orthologs either lack the supporting hydrogen-bond network provided by Y457, instead harbouring non-polar aromatic amino acids (e.g., F457) or have Y457 adopting a different rotamer where it cannot participate in the same H-bond network. This is consistent with previous work that highlighted that, even in *Lcα*E7, the G137D mutation only transiently sampled catalytically productive conformations (22). Thus, if the productive conformation of Asp137 cannot be sampled due to the surrounding amino acids, it would not be catalytically beneficial. These structural discrepancies also help explain why G137D causes a substantial *K*_m_ increase, given it simultaneously disrupts the active site and binding of the OP. Additionally, loop regions surrounding the active site differ significantly across orthologs, affecting substrate binding and how the Asp137 mutants will interact with OPs (**Figure 2D**). Taken together, these analyses highlight that no single mutation exists in isolation; rather, the impact of G137D is shaped by the collective geometry and chemistry of the active site, emphasizing the structural basis for epistasis and historical contingency in this example of neofunctionalization.

**Figure 2:**
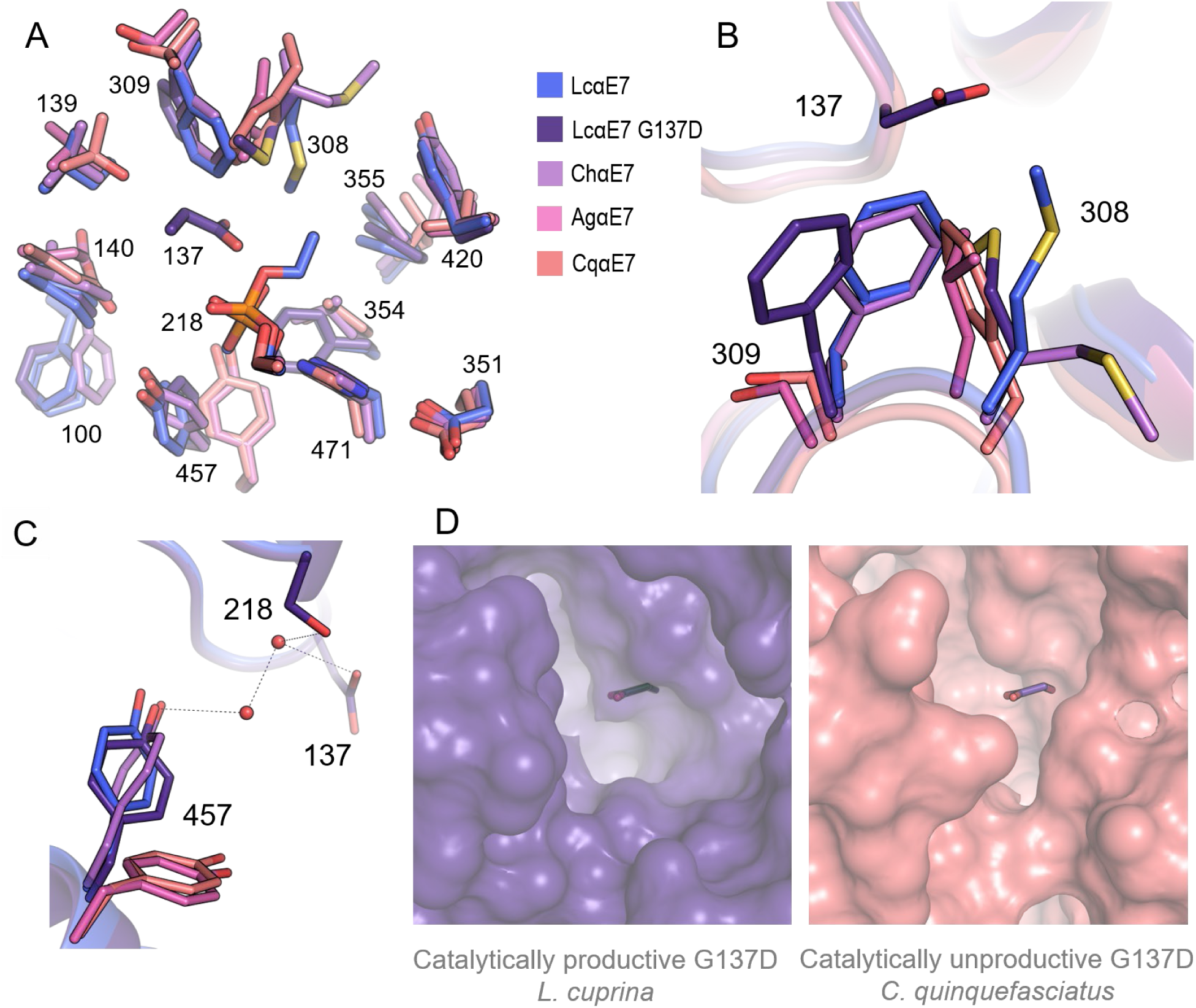
Structural basis for the sequence context dependence of the G137D mutation. Phosphorylated wild type *Lcα*E7 is shown in blue (PDB ID: 5IVI), *Lcα*E7 G137D is shown in dark purple (PDB ID: 5C8V), phosphorylated *Chα*E7 (PDB ID: 9D1J) is shown in light purple, *Agα*E7 (PDB DI: 9D1K) is shown in pink, and *Cqα*E7 (PDB ID: 5W1U) is shown in coral. **A)** Overlay of the active sites. Side chains of key residues are shown in sticks. **B)** Residues Phe309 and Met308 in *Lcα*E7 constrain Asp137 in a catalytically favourable conformation. **C)** A unique conformation of Tyr457 in *Lcα*E7 enables an energetically-favourable water bonding network within the active site. **D)** Comparison of the active site surface of *Lcα*E7 *vs. Cqα*E7 shows a more open active site for *Lcα*E7.

### Directed evolution of *Lcα*E7 for increased OP hydrolase activity

To investigate how *L. cuprina α*E7 (*Lcα*E7) could continue to evolve to enhance organophosphate (OP) hydrolysis further, we used directed evolution. Starting with a wild-type *Lcα*E7 plasmid, we employed error-prone PCR (epPCR) to introduce random mutations (1-3 per gene), generating variant libraries. Each library, transformed into *E. coli*, yielded ∼2,000 colonies per round, all screened for improved hydrolysis of the fluorogenic analogue diethylumbelliferyl phosphate (DEUP) in 96 well plates. The best 5-10 variants from each round were pooled for the subsequent round. After isolating the best variant from each of the nine rounds (R1–R9) (**Figure 3A,B,C**), we investigated the basis for how these mutations improve OP hydrolysis using stopped-flow fluorescence to track changes in the phosphorylation (*k*_3_) and dephosphorylation rates (*k*_5_) (**Figure 3D, Supplementary Table 3**). The pre-steady state burst was evident in early rounds (R1-R6), with clear biphasic (burst followed by steady-state) kinetics. However, in later rounds (R7-R9), once the rate of dephosphorylation (*k*_5_) exceeded 0.06 s^-1^ and the burst rate decreased, the reaction resembled classical Michaelis– Menten behaviour without an observable burst phase. These results show progressive improvement in the steady state turnover rate for *Lcα*E7, eventually reaching rates 1100-fold higher than wild-type, although the largest increase was observed in R1 (G137D; 18-fold), with later mutations conferring smaller (1.4 to 9-fold) relative improvements. Notably, the Michaelis constant (*K*_m_) for DEUP hydrolysis increased 33-fold across the evolutionary trajectory, with the R9 variant exhibiting a *K*_m_ (49 µM) substantially higher than R1 (<1.5 µM). By the ninth round, variant *Lcα*E7-9 contained 11 mutations, with most of the later mutations being located in remote regions but still influencing performance (**Figure 3C**). This outward radiation of mutations along directed evolution trajectories is often seen and is consistent with a model where initial mutations affect active site chemistry (G137D), followed by mutations that stabilize the new active site (e.g. M308V), followed by mutations that fine tune protein conformational changes (36).

**Figure 3:**
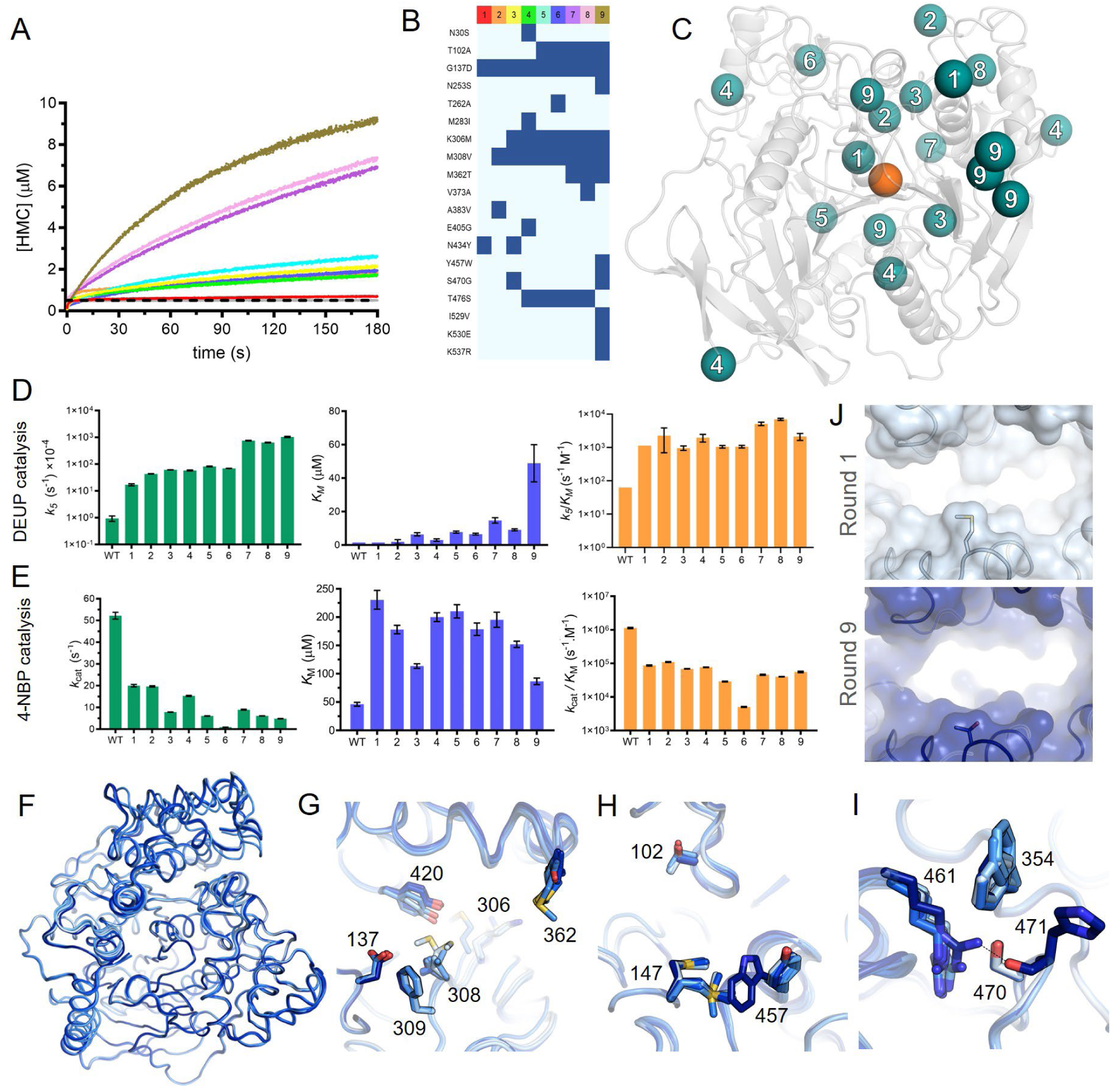
Directed evolution of *L. cuprina α*E7. **A)** Representative stopped-flow kinetic data for OP-hydrolysis by *Lcα*E7 WT and evolved variants. Colors as indicated in panel B. The concentration of product HMC (7-hydroxy-4-methyl coumarin) is indicated. **B)** Amino acid substitution in the directed evolution rounds relative to WT. **C)** Mutations relative to WT are highlighted in teal spheres, numbered by round, on the *Lcα*E7 backbone (grey) and were distributed throughout the enzyme. The catalytic serine is showed in an orange sphere **D)** Kinetic parameters for OP hydrolysis, values are mean ± standard error (n=4). **E)** Kinetic parameters for ester hydrolysis values are mean ± standard error (n=3). **F)** Alignment of the R1-R9 crystal structures, colored from R1 (light blue) to R9 (dark blue). **G)** Mutations along the evolutionary trajectory, such as Met308Val, Lys306Met, and Met362Thr, reconfigured the interactions near the active site. **H)** Tyr457Trp mutation in R9 caused a displacement in the flexible residue Met147, opening the binding pocket. **I)** The Ser470Gly mutation in R3 and R9 enabled the insertion of Arg461 into the active site, forming a new hydrogen bond with the backbone carbonyl of His471. **J)** Present from R7-R9, the Met362Thr mutation (shown in sticks) created a wider opening to the binding pocket of the enzyme.

Because *Lcα*E7 functions as a general carboxylesterase, we next tested how these evolutionary changes altered its native esterase activity using 4-nitrophenyl butyrate (**Figure 3E, Supplementary Table 4**). Even the initial G137D variant (R1) displayed a notable drop in *k*_cat_ for 4-NPB, implying that D137 disrupts the enzyme’s optimal geometry for ester substrates. As additional mutations accumulated, some later variants retained only less than 10% of the wild-type esterase *k*_cat_, illustrating a growing trade-off between new and ancestral functions. Consequently, *Lcα*E7 became progressively specialized for OP detoxification while sacrificing esterase efficiency. From an evolutionary standpoint, this highlights how a single-site change like G137D could initiate a cascade of mutations that drive enzyme adaptation in pesticide-laden environments, where enhanced detoxification often outweighs modest losses in native activity. However, there is no evidence of this occurring beyond compensatory intergenic epistatic modifiers mutations or duplications to restore enough esterase function to support essential physiological roles (37, 38). In this manner, our directed evolution study illustrates a hypothetical *in vivo* process, revealing how *Lcα*E7’s capacity for mutation and rearrangement can, in principle, yield a highly effective OP hydrolase.

A key question arising from our directed evolution of *L*. *cuprina α*E7 (*Lcα*E7) is how incremental mutations collectively reshape the enzyme’s active site and surrounding regions to favour organophosphate (OP) hydrolysis. To address this, we solved high resolution crystal structures of each of the evolution rounds (**Figure 3F, Supplementary Table 1**). The G137D mutation is the mechanism changing “linchpin” mutation, which we have previously shown initially exists in a disordered state, sampling both productive geometries where it allows for substrate binding and orients/activates a water molecule for hydrolysis of the phosphorylated intermediate, and a non-productive conformation that blocks the active site (22). Subsequent mutations along the directed evolution trajectory collectively refined the active site architecture to overcome these limitations, leading to further substantial increases in *k*_5_. Mutations such as Met308Val, Lys306Met, and Met362Thr reconfigured the interaction between this region and the Asp137/Phe309 pair (**Figure 3G**). The reduction in steric bulk at position 308 and the disruption of a salt-bridge at Lys306 facilitated a shift in the loop, promoting a more compact state stabilized by a persistent hydrogen bond between Tyr420 and the Met306 backbone carbonyl. This conformational adjustment of the loop is proposed to subtly reorient the Asp137/Phe309 pair, favoring a more open and catalytically competent conformation that facilitates efficient dephosphorylation. Concurrently, mutations such as Thr102Ala and Tyr457Trp may influence active site dynamics: while Thr102Ala’s precise mechanism remains subtle, Tyr457Trp, accommodated by the flexibility of Met147, induces an opening of the binding pocket, which may destabilize the phosphorylated intermediate and accelerate its release (**Figure 3H**). Furthermore, the Ser470Gly mutation enabled the insertion of Arg461 into the active site, forming a new hydrogen bond with the backbone carbonyl of the catalytic histidine (**Figure 3I**). This new catalytic motif, involving Arg461 and a concomitant rotation of Phe354 to alleviate steric clashes, likely further contributes to the enhanced dephosphorylation rate. Additionally, R7-R9 displayed a larger binding site opening (**Figure 3J**) due to the Met362Thr mutation. This observation is consistent with the increased *k*_5_ in R7-9 for hydrolysis of the bulkier substrate DEUP. Collectively, these diverse, often remote, mutations act synergistically to fine-tune the enzyme’s conformational landscape, reducing the energetic barriers for the dephosphorylation step and dramatically increasing *k*_5_. While unique, the trend is one that is seen repeatedly in enzyme evolution; an initial mutation perturbs the active site chemistry but is initially imperfect owing to conformational sampling of non-productive states; subsequent, increasingly remote mutations then refine the conformational landscape to optimize the preorganization of the active site for its new chemistry/substrate, often in overlapping partially redundant ways.

## *In vivo* resistance conferred by laboratory evolved *α*E7

To investigate the *in vivo* relevance of enhanced organophosphate (OP) detoxification, we generated transgenic Drosophila expressing wild type *α*E7 and evolved variants under the control of a promoter that produced elevated levels of expression of *α*E7 in three key metabolic tissues: the midgut, Malpighian tubules and fat body (39). Following confirmation that each of the transgenic lines were overexpressing their respective allele of the *α*E7 gene (**Supplementary Figure 3**), we used toxicology screens to determine their resistance levels to the structurally analogous diethyl organophosphate insecticides paraoxon and diazinon (**Figure 4A**). It is worth noting that the lethal concentration at which 50% of flies die (LC_50_) values that are measured are a consequence of the inhibition of the target AChE, not a direct assay of the detoxification enzyme *Lcα*E7.

**Figure 4:**
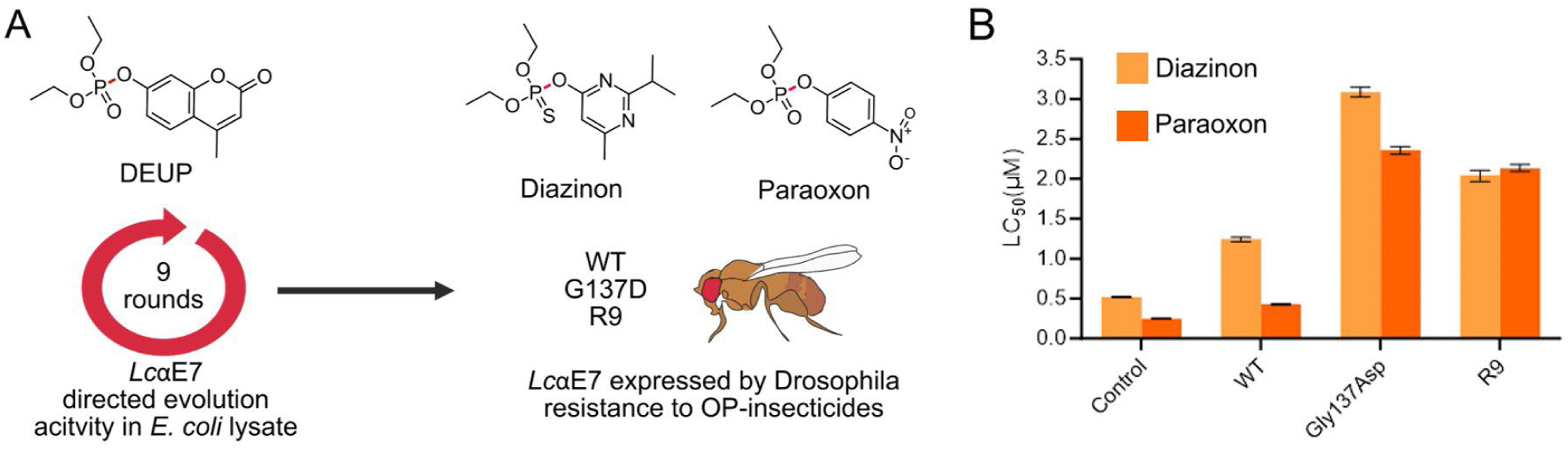
Expression of *Lcα*E7 variants in *Drosophila* confers insecticide resistance. **A)** Nine rounds of directed evolution resulted in *Lcα*E7 variants with increased rates of OP-hydrolysis. Three *Lcα*E7 genes (wild-type, G137D and R9) were introduced into the genome of *Drosophila*. Fluorometric substrate DEUP, and insecticides Diazinon and Paraoxon are shown with the scissile bond indicated in red. **B)** Diazinon and Paraoxon LC_50_ (concentration that kills 50 % of flies) were measured. Control is the *Drosophila* background line without a transgene, wild-type is wild-type *Lcα*E7. Values are mean, bars show upper and lower limit. (n=1250 larvae)

Our data reveal a significant disconnect between the *in vitro* catalytic performance of the evolved *Lcα*E7 variants and the *in vivo* resistance they confer when introduced into *D. melanogaster*. Although the R9 enzyme, the final product of our directed evolution, displayed a more than 60-fold higher turnover number (*k*_5_) against diethyl OP substrates *in vitro* (**Figure 3D**) compared to the G137D variant, it conferred reduced resistance to OPs *in vivo* (**Figure 4B, Supplementary Tables 5-6**). For example, with diazinon, the LC_50_ for flies expressing G137D was approximately 3.0 µM, whereas for R9 it was approximately 2.0 µM, indicating G137D conferred greater protection. Similarly, for paraoxon, the LC_50_ for G137D was slightly higher (≈2.4 µM) compared to the transgenic fly harbouring R9 (≈2.1 µM). This counterintuitive outcome highlights that an enhanced catalytic rate (*k*_5_) and greater turnover is insufficient to guarantee superior *in vivo* protection if it is accompanied by a substantial loss of substrate affinity, which kinetically manifests as an increased Michaelis constant (*K*_m_), which in the case of the G137D *vs.* R9 comparison, *K*_m_ increased from 1.5 µM to approximately 50 µM (**Figure 3D**).

To understand this phenomenon, we must consider the kinetic parameters in the physiological context of OP exposure. The target enzyme, *D. melanogaster* AChE, possesses an inhibition constant (K_i_) for typical OPs (like paraoxon and diazinon) in the order of 1.5 µM, which is consistent with the observed LC_50_ values (e.g., ≈1 µM for wild type flies (40). At these low micromolar concentrations, the G137D variant, with its *K*_m_ of approximately 3 µM, is well-suited to bind OPs. This *K*_m_ value is in the same range as the toxic OP concentrations and the sensitivity of the target AChE (*K*_i_≈1.5 µM), meaning G137D can effectively capture a good portion of the insecticide. In contrast, the R9 variant has a much higher *K*_m_ of approximately 50 µM. This means that at the same low-micromolar OP concentrations (around 2.5 µM), R9 binds the insecticide much less effectively. While G137D would bind a substantial fraction of the OP (roughly 45% of its enzyme molecules would be occupied at an OP concentration of 2.5 µM), R9 would bind very little (only about 5% of its enzyme molecules would be occupied at the same OP concentration). This suggests that G137D would be much more likely to bind an OP molecule compared to R9 at these low concentrations. Thus, even though R9 is much faster at breaking down the OP once it binds (over 100-fold higher *k*_5_), the improved catalytic efficiency per encounter is rendered less effective *in vivo* because R9 has fewer encounters with the OP molecules at the relevant toxicological concentrations and more insecticide molecules bypass R9 and reach the sensitive AChE target. Clearly, turnover (*k*_5_) is beneficial, as we observe a significant increase in LC_50_ for both G137D and R9, relative to WT *Lcα*E7, even though WT has sub-micromolar *K*_m_, which suggests that the wild-type is rapidly saturated and unable to provide a detoxification benefit above a certain level. These observations underscore the delicate balance that must be maintained between substrate affinity and binding interactions in conferring robust *in vivo* resistance and suggest that any natural evolution for increased resistance will need to increase turnover, while maintaining high affinity.

## Discussion

The ability of certain dipteran insects to acquire organophosphate (OP) resistance via a single Gly→Asp active site mutation in alpha-carboxylesterase 7 (*α*E7) exemplifies how historical contingency and epistasis determine and constrain molecular evolutionary trajectories. In the case of *L. cuprina*, the enzyme’s active site architecture and surrounding residues, particularly Met308, Phe309, and Tyr457, provide a uniquely accommodating environment for Asp137 to allow it to adopt a catalytically productive conformation, allowing dephosphorylation of the phospho-serine intermediate. In environments where OPs are present, this local structural compatibility allows G137D to act as a beneficial change rather than a neutral or deleterious substitution. Conversely, for *α*E7 proteins in species, such as *C. quinquefasciatus*, that lack this supportive network, G137D offers little or no catalytic advantage. This type of conformational epistasis, where the effect of a sequence change depends on the conformational sampling of the amino acid in space, underscores how historical sequence changes can either potentiate or preclude the success of G137D, shaping which insect populations can exploit this solution. Indeed, field observations reveal that OP selection pressure has driven G137D alleles to near fixation in *L. cuprina* and a handful of other flies, while many equally exposed insect species rely instead on alternative mechanisms such as gene amplification, or target site (AChE) modifications (41). This divergence highlights the importance of structural pre-adaptations, whether in key loop regions, hydrogen-bonding networks, or backbone conformations, that can potentiate a protein such that an otherwise deleterious mutation becomes beneficial. Thus, the success of G137D-based OP resistance in certain dipteran lineages can be explained by the interplay of contingency, which laid down the permissive sequence context, and epistasis, which dictates how well new mutations integrate into an existing protein scaffold. This combination of historical and structural factors ultimately determines which species acquire potent catalytic detoxification and which remain locked into less efficient resistance strategies. Similar observations have been made in antibiotic resistance proteins in model systems (27, 31, 33), but this is a relatively rare real-world, eukaryotic example.

Our directed evolution experiments demonstrate that a single function-altering active site mutation like G137D can be the starting point for more extensive protein adaptation under strong selection. By screening mutant libraries of *L. cuprina α*E7 at relatively high OP concentrations, we effectively biased the evolutionary trajectory toward variants that excel at regenerating the active site once phosphorylated, i.e., variants with higher dephosphorylation rates. We observe a common pattern in laboratory-evolved enzymes (R1 through R9) where additional substitutions accumulate radially out from the initial G137D mutation. The mechanistic basis for their beneficial effects appears to involve optimizing the conformational sampling of catalytic residues by stabilizing crucial interactions. Structural analysis reveals that these changes in peripheral loops and helices remodel the free-energy landscape of the protein, such that phospho-serine intermediates are broken down more efficiently, either by reducing stabilization of the intermediate or optimizing the orientation of the general base G137D to generate the attacking nucleophile. However, this focus on improving turnover at high substrate concentrations also exerts a strong penalty on the enzyme’s substrate affinity. Indeed, the evolved variants consistently exhibit higher *K*_m_ values, indicating reduced OP binding affinity; an unsurprising trade-off given that many beneficial mutations can expand or reorganize the active site, and stabilizing the transition state for turnover often compromises substrate capture. Moreover, these gains in OP hydrolysis are tied to substantial losses in native esterase function (4-nitrophenyl butyrate hydrolysis), even in the earliest round containing G137D. This trajectory reflects a strong positive trade-off: each new mutation propels *α*E7 further from its ancestral function to intensify catalytic throughput against OPs, but at the cost of diminishing native substrate capacity. In nature, mutations in other genes or duplication of the wild type gene has compensated for the loss of the ancestral function (10, 42).

Although the directed evolution approach yielded *α*E7 variants with dramatically higher catalytic activity *in vitro*, introducing these variants into *D. melanogaster* made it clear that the enhanced turnover does not necessarily equate to superior *in vivo* protection. In fact, the R9 mutant, which showed a remarkable increase in dephosphorylation rate *in vitro*, offered less protection than the simpler G137D variant when flies were exposed to physiologically relevant (low) OP concentrations. This underscores that high substrate turnover is only beneficial if the enzyme can bind and sequester the toxin at sub-saturating levels; a critical requirement given the extreme toxicity of OPs. Because the R9 mutant’s improvements in dephosphorylation were accompanied by a loss of affinity (increased *K*_m_), any catalytic advantage is lost because the OPs can evade R9 to bind preferentially to the target AChE. Ecologically and evolutionarily, this points to the primary role of *α*E7 as an “OP sponge” in real-world scenarios, where capturing low levels of insecticide is a pre-requisite over substrate turnover, i.e., turnover (dephosphorylation) is valuable for regenerating the unbound enzyme pool, but requires high affinity OP binding. From an adaptive standpoint, it is therefore not surprising that no additional high-turnover G137D modifications have become widespread in field populations. Instead, insects have primarily evolved epistatic changes that restore partial esterase function or enhance protein stability, including gene duplications that alleviate the cost of lost native activity. Hence, field-level selection strongly favours an optimal balance of sequestration and moderate turnover over the purely turnover-centric optimization we observed in our *in vitro* experiments.

Taken together, these findings illustrate how evolutionary pathways to OP resistance rely on the interplay of historical contingency, epistasis, and balancing selection pressures. First, only esterases that have a comparable substrate affinity for OPs than the target AChE are candidates (43). Second, only dipteran lineages already primed by specific structural or sequence features can readily exploit G137D to become potent OP hydrolases. Third, directed evolution in the laboratory underscores how an initial advantageous mutation can result in a cascade of secondary substitutions that raise dephosphorylation rates but that the requirement to simultaneously increase turnover while retaining high affinity is a challenging problem to overcome. These results also highlight that translating *in vitro* enzymatic enhancements into *in vivo* resistance of generalized physiological conclusions depends critically on many factors; in this case, whether those variants bind toxins effectively at the low concentrations encountered in the environment. It suggests caution when interpreting the physiological implications of these studies. Our transgenic fly studies confirm that maximizing turnover without preserving binding affinity fails to deliver improved protection. Thus, while our work explores the potential to reconfigure *α*E7 through laboratory manipulation, it also highlights the context-dependent nature of enzyme adaptation. Insects have large multigene families of genes that, in theory, provide many options in defending them against insecticides via enhanced metabolism (44). In reality, the number of real options are narrowed down by many factors including the necessity for appropriate patterns of gene expression (45). This study has shown that even for a gene that is appropriately expressed, beneficial mutations must maintain a balance between catalytic turnover and substrate affinity. By delineating these trade-offs, we advance our understanding of enzyme evolution and provide insight into how insects respond, or fail to respond, when challenged by potent chemical threats.

## Materials and Methods

### Protein expression and purification

All *α*E7 variants were expressed from the pETMCSIII vector with an N-terminal His-tag. Protein was expressed in BL21(DE3) cells (NEB, C2527) using lysogeny broth (LB) (10 g/l NaCl, 10 g/l tryptone, 5 g/l yeast extract) supplemented with 100 μg/mL ampicillin. After growth at 24°C for 20 hours, cells were pelleted by centrifugation (5000 g, 14 min, 4°C) and then resuspended in binding buffer (300 mM NaCl, 20 mM imidazole and 50 mM HEPES pH 7.5) supplemented with 5 units/mL turbonuclease (Sigma, T4330). Approximately 5 mL of binding buffer was used to resuspend each gram of wet cell pellet. Resuspended cells were supplemented with 0.2% Triton X100, vortexed thoroughly and lysed via sonication. Lysate was clarified by centrifugation (13,500 g, 45 minutes), and the soluble fraction was applied to a 5 mL HisTrap HP nickel affinity column (GE Healthcare). The column was washed with 5 column volumes binding buffer followed by 5 column volumes 7% elution buffer (300 mM NaCl, 300 mM imidazole, 50 mM HEPES pH 7.5). The bound protein was eluted with 100% elution buffer and the single 5 mL fraction containing *α*E7 was subjected to size exclusion chromatography with a Superdex 26/600 200 pg column (GE Healthcare) with 150 mM NaCl, and 20 mM HEPES pH 7.5. Protein was quantified based on absorbance at 280 nm using an extinction coefficient calculated with the Protparam online tool (https://web.expasy.org/protparam/). Typical yields were 10-20 mg per litre of media. There was soluble expression of the recombinant carboxylesterases from each of the directed evolution intermediates and the orthologs except for *Hiα*E7 which was largely insoluble and was subsequently not analyzed in further detail.

### Esterase activity assays

Esterase activity was determined with the model ester 4-nitrophenyl butyrate (4-NPB; Sigma, N9876). Reactions consisted of 148 μL assay buffer (150 mM NaCl, 20 mM HEPES-NaOH pH 7.5), 2 μL substrate (prepared in 100% DMSO) and 50 μL enzyme (prepared in assay buffer supplemented with 1 mg/mL bovine serum albumin (BSA; Sigma, A7030)). Enzyme solutions were stored on ice until immediately prior to use. After the addition of enzyme, the reactions were mixed thoroughly and formation of the 4-nitrophenol product of hydrolysis was monitored at 405 nm at 24°C for 2 minutes using a microplate spectrometer (Biotek, Epoch 2). Initial velocities were corrected for nonenzymatic hydrolysis, converted into μM/second using a standard curve of 4-nitrophenol, and the parameters Vmax and *K*_m_ were determined by fitting the velocities to the Michaelis-Menten equation using non-linear regression in GraphPad Prism 7.

### OP-hydrolase activity assays

OP-hydrolase activity was determined with diethylumbelliferyl phosphate (DEUP; Sigma, D7692) in assay buffer (150 mM NaCl, 20 mM HEPES-NaOH pH 7.5) as previously described (22). In brief, solutions containing DEUP were mixed with enzyme (final concentration 0.75 µM) using a RX2000 rapid mixing stopped-flow unit fitted with a RX/DA pneumatic drive (Applied Photophysics, UK). The hydrolysis of DEUP was monitored fluorometrically (excitation at 330 nm and emission at 450 nm) with a Cary Eclipse Fluorescence spectrophotometer (Agilent Technologies, USA).

### Crystallization and structure determination

Crystallization of *α*E7 was achieved by reductive lysine methylation(46). Proteins were first purified by nickel affinity chromatography and size exclusion chromatography as described previously, except that the size exclusion step was performed using 150 mM NaCl, 50 mM HEPES pH 7.5. On the same day as purification, the methylation was performed by adding 20 μL 1 M dimethylamine borane complex (prepared in water) (Sigma, 180238) and 40 μL 1 M formaldehyde (prepared in water) (Sigma, 252549) per 1 mL protein solution (previously diluted to 0.8 mg/mL with size exclusion buffer). The reaction was assembled on ice in a 50 mL tube and was typically performed with 20 mg of protein at a time. After two hours of slow agitation at 4°C, the first step was repeated, and the solution returned to 4°C for an additional 2 hours. In the final step of the methylation reaction, 10 μL 1 M dimethylamine borane complex solution was added per mL protein, and the mixture was left at 4°C for 12 hours. To ensure all formaldehyde was consumed, and to reduce any intermolecular disulfides, 125 μL 1 M glycine (prepared in water) and 125 μL 0.05 M dithiothreitol (prepared in water) was added per mL protein solution, and the reaction was agitated slowly at 4°C for an additional two hours. The sample was concentrated using a 30 kDa molecular weight cut-off spin concentrator (Amicon) and subjected to size exclusion chromatography using 150 mM NaCl, 20 mM HEPES pH 7.5. The final size exclusion step removed excess reagents and soluble protein aggregate. Immediately after size-exclusion, protein was diluted 1:2 with water, and concentrated for crystal screens.

Crystals of WT *α*E7 were grown at 19°C using hanging drop vapor diffusion with a reservoir containing 500 μL 200 mM BisTris pH 6.5, 19-21% polyethylene glycol (PEG) 3350 and 6% methyl-2,4-pentanediol (MPD). Drops consisted of 4 μL protein (10 mg/mL) and 2 μL reservoir solution. Crystals with dimensions of approximately 200 × 50 × 50 μm grew overnight. To crystalize the evolved variants, and to obtain large crystals of WT *α*E7, microseeding was performed. Wild type crystals were crushed on a glass coverslip using a glass rod and then transferred to a 1.5 mL tube containing 500 μL freshly prepared reservoir solution matching the crystallization condition. The seed mixture was vortexed, and serial 1:10 dilutions were prepared. Optimization of crystal size and growth was typically achieved by varying PEG concentrations from 8 to 13% (with 200 mM BisTris pH 6.5 and 6% MPD) and varying seed stock concentration from 1:10 to 1:10000. Hanging drops consisted of 2 μL protein, 1.6 μL reservoir and 0.4 μL seeding solution. Crystals grew within 1-3 days and seeds stocks were used reproducibly for up to 2 weeks when stored at 19°C.

Crystals were susceptible to cracking when removed from the drops. To avoid this and to achieve cryoprotection, a volume of cryoprotectant (35% PEG 3350, 10% MPD, 200 mM BisTris pH 6.5) equal to the drop volume was pipetted directly onto the drops. Crystals were subsequently retrieved from the original drop through the cryoprotectant solution and then flash cooled in liquid nitrogen. *Agα*E7 crystals were obtained using 1 µl 13 mg/mL protein solution + 1 µl well solution equilibrated against a well solution of 24 % PEG 4000, 0.2 M MgCl_2_, 0.1M Tris pH 8.1. *Chα*E7 crystals were obtained using 20-25 % PEG3350, 0.1M MES pH 6.5. To capture the phosphorylated *Chα*E7 structure, cryoprotectant solution was prepared with DEUP at 1 mM (with DMSO at 1%), the cryoprotectant was pipetted onto the crystallization drops, and the crystals were fished into the solution and allowed to soak for < 5 minutes, before being flash frozen.

Data for all structures were collected on the MX2 beamline at the Australian Synchrotron(47), and diffraction images were processed using XDS(48) in space group P 2_1_ 2_1_ 2_1_ for all directed evolution and *Chα*E7 structures and P 1 2_1_ 1 for the *Agα*E7 structure. For all structures, data were truncated using Aimless(49) until statistics were of suitable quality in the outer shell. The structures were solved by molecular replacement (Phaser MR, CCP4(50)) using PDB ID 5C8V as the search model for *Lcα*E7 structures, and AlphaFold2 structures for *Agα*E7 and *Chα*E7. Refinement was carried out in Phenix.refine(51). Multiple rounds of refinement and model building in Coot(52) were carried out until R-factors converged and the electron density was modeled as best as possible. MolProbity(53) was used for structure validation and Pymol 2.0 was used for figure generation. Structures have been deposited in the Protein Data Bank under IDs 9D1J, 9D1K, 9D1L, 9D1M, 9D1N, 9D1O, 9D1P, 9D1Q, 9D1R, 9D1S, 9D1T.

### Transgenic expression of Lcα*E7* genes in D. melanogaster

The sequences of *Lcα*E7 genes (WT and R9) were codon optimised for expression in *D. melanogaster* with Thermo Fisher’s Gene Optimiser (**Supplementary Table 7**). Genes were designed with flanking 5’ and 3’ *Eco*R1 sites. Sequences were obtained from Invitrogen and cloned into the pETMCSI vector using Gibson assembly. The G137D construct was generated by introducing the G137D mutation into the *Lcα*E7 WT gene using primers through Gibson assembly. Each plasmid construct was digested using *Eco*R1 restriction enzyme (Promega), ligated into the pUAST-attB plasmid(54), and transformed into silver α-selected competent *E. coli* cells (Bioline). Plasmid DNA was prepared using the Nano-plasmid mini-extraction kit (Bioneer) and then sequenced. Constructs with the correct DNA sequence were microinjected into pre-blastoderm *D. melanogaster* embryos containing an attP40 landing site and expressing phiC31. The *white* marker allowed transformants carrying an integrated construct to be identified. Homozygous transformed lines were made and the successful integration of the pUAST-attB construct was confirmed through sequencing. To generate a control line with the same genetic background (Control) an empty pUAST-attB vector was injected and incorporated into the genome. The GAL/UAS system was used to overexpress each *Lcα*E7 gene(55), with males of each UAS-*Lcα*E7 line crossed to females of the HR-GAL4 driver line. This resulted in offspring overexpressing the *Lcα*E7 gene in the midgut, Malpighian tubules and fat body(39). All fly lines used in this study are listed in **Supplementary Table 8**.

Expression levels of *Lcα*E7 gene and the variant alleles were quantified using reverse transcription polymerase chain reaction (RT-PCR). For each line, RNA was extracted from three biological replicates of 25 third instar larvae using TRIzol reagent (Bioline). Each RNA sample was treated with RNase-free DNase (Promega). Reverse transcription was then performed on 1 μg of each RNA sample in a 20 μL reaction using GoScript (Promega) and oligo(dT)20 primer, following the manufacturer’s instructions. RT-PCR was conducted on 2 μL of a 1 in 10 dilution of cDNA using SYBR-Green (QIAGEN) and the CFX384 Touch RT-PCR detection system (BioRad). Primers used are listed in **Supplementary Table 7**. PCR conditions were 95°C for 5 mins for enzyme activation, followed by 40 cycles of 95°C for 10 secs and 60°C for 30 secs. Relative expression of each gene was measured in reference to the housekeeping gene *Rpl11* and *CG13220*. To compare the expression levels of each transgenic line, a One-way ANOVA test with post-hoc Tukey’s HSD (p≤0.05) was performed.

### Evaluating resistance levels in first instar larvae

Resistance levels were measured using standard first instar toxicology assays as described previously(56). First instar larvae of each *Lcα*E7 line were exposed to the OP insecticides diazinon (Mallinckrodt Veterinary) and paraoxon (Sigma-Aldrich). Insecticides were diluted in dimethyl sulfoxide (DMSO) and mixed with fly food media at the appropriate concentrations. Flies from each *Lcα*E7 overexpression line were allowed to lay on apple juice plates partially covered in yeast paste. Laid eggs were collected daily, washed, and left to hatch on new plates overnight. For each of the 5 doses used, 5 replicates of 50 first instar larvae were transferred onto vials of insecticide-dosed maize meal medium (1250 larvae screened for each fly line). Vials were kept in the dark at 25°C and scored for eclosion up to 16 days after the larval transfer. Any fully eclosed adults, even if dead at the time of scoring, were considered to have survived the assay. Results were corrected for control mortality using Abbott’s correction and a Probit analysis was used to calculate the LC_50_(57, 58). All analysis was done in R software using a modified script from an analysis package obtained from GitHub (https://github.com/shanedenecke/insect.toxicology). Graphs and figures were generated using GraphPad Prism 8 (GraphPad Software, CA, USA).

## Acknowledgments

This work was supported by the Australian Research Council Centre of Excellence for Innovations in Protein and Peptide Science (CE200100012) and the Australian Research Council Centre of Excellence in Synthetic Biology (CE200100029). C.J.J. was supported by an Australian Research Council Future Fellowship. P.D.M. was the recipient of a John Stocker Postdoctoral Fellowship from the Science and Industry Endowment Fund (Australia). This research was undertaken in part using the MX2 beamline at the Australian Synchrotron, part of ANSTO, and made use of the Australian Cancer Research Foundation (ACRF) detector. We dedicate this work to the memory of Prof. Dan S. Tawfik, whose interest, insight and guidance shaped this study over many years.

## Author contributions

C.J.J., P.D.M., P.B., and J.O. designed research; A.G., M.E., P.D.M. and N.F performed research; J-W. L., performed directed evolution; R.L.F., A.G., P.D.M., and C.J.J performed data analysis; P.D.C., N.F., and D.H. crystallised proteins and collected crystallographic data; R.L.F and C.J.J. performed crystallographic data processing and analysis; R.L.F., A.G., P.D.M., and C.J.J. prepared the manuscript with contributions from all authors.

## Supporting information

**Supplementary figure 1:**
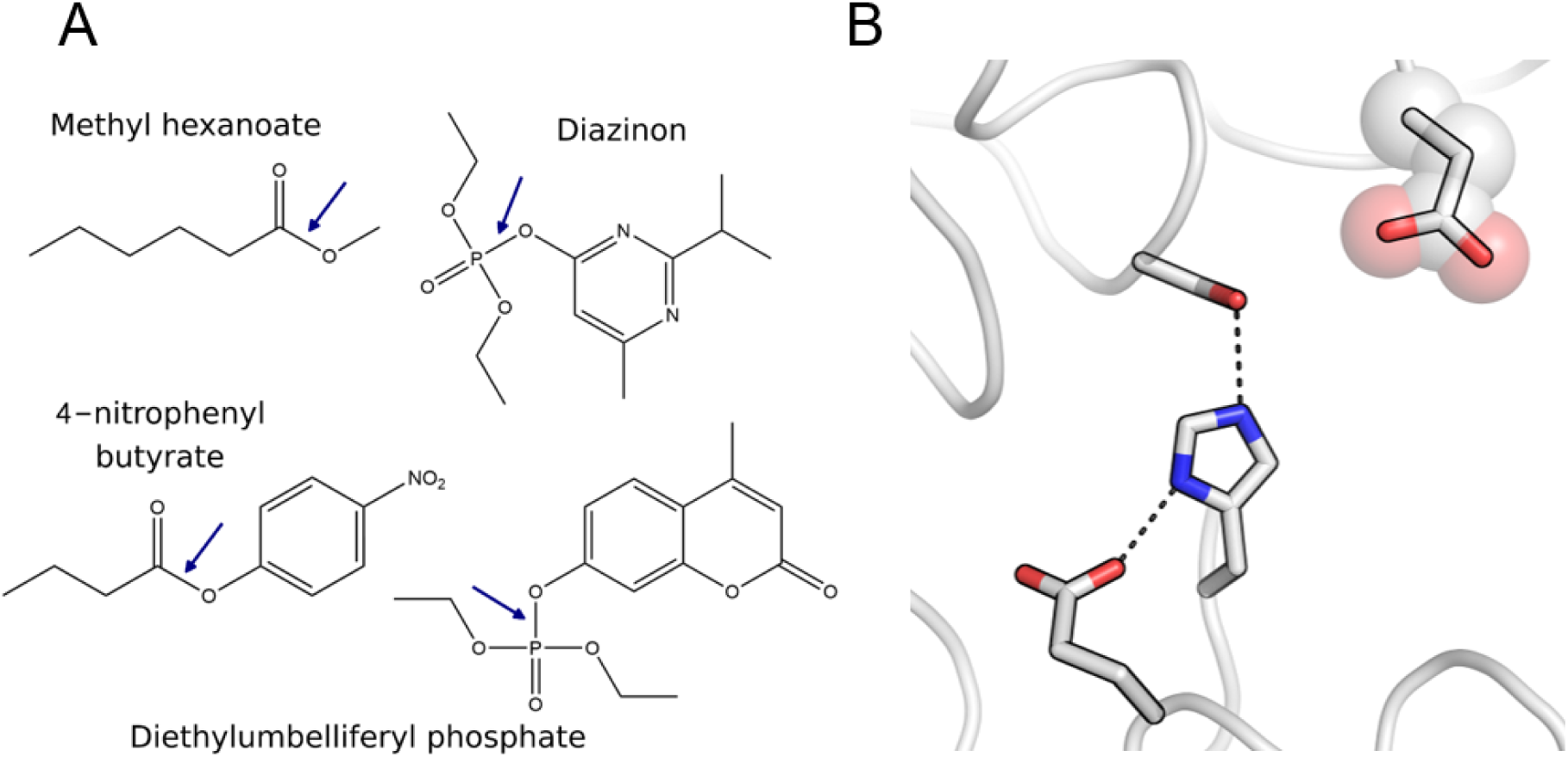
**(A)** a substrate of *L. cuprina α*E7 (Methyl hexanoate), the diethyl-OP insecticide (Diazinon), the chromogenic ester-substrate (4-nitrophenbyl butyrate) and the fluorogenic OP-substrate (Diethylumbelliferyl phosphate). **(B)** Catalytic triad (Ser218, Glu351 His471) and Asp137 in *Lcα*E7 (PDB 5C8V).

**Supplementary figure 2:**
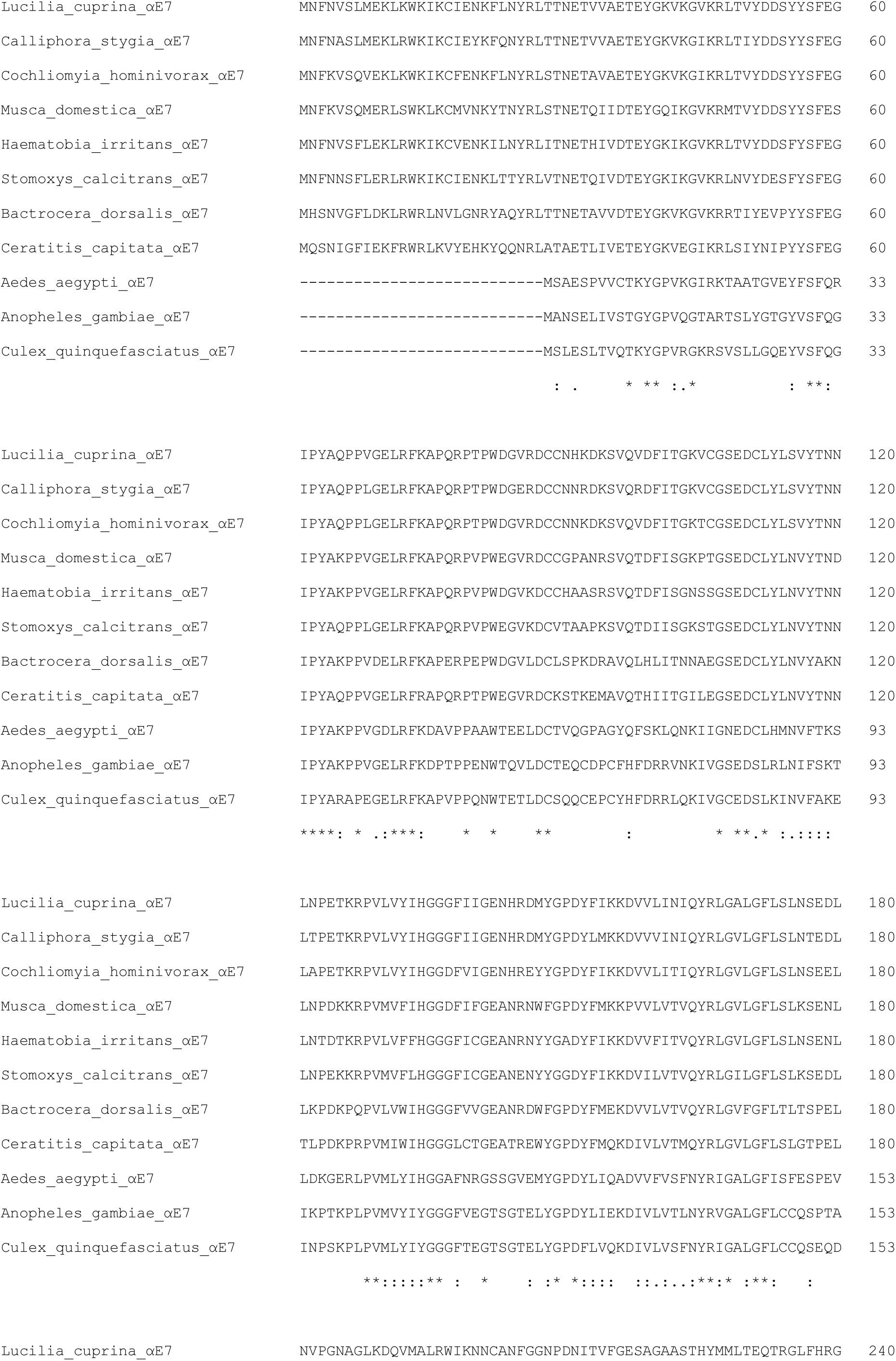
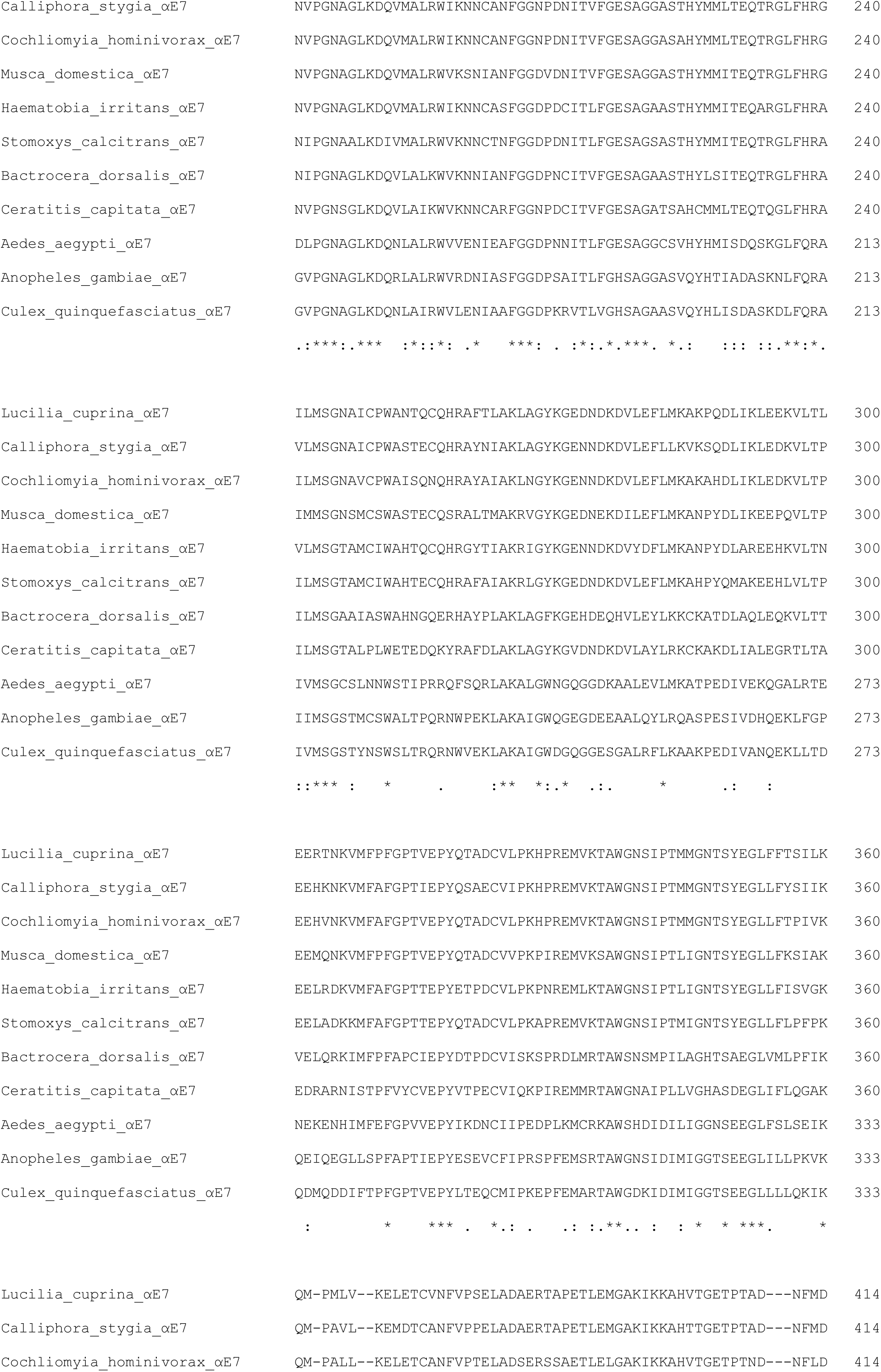
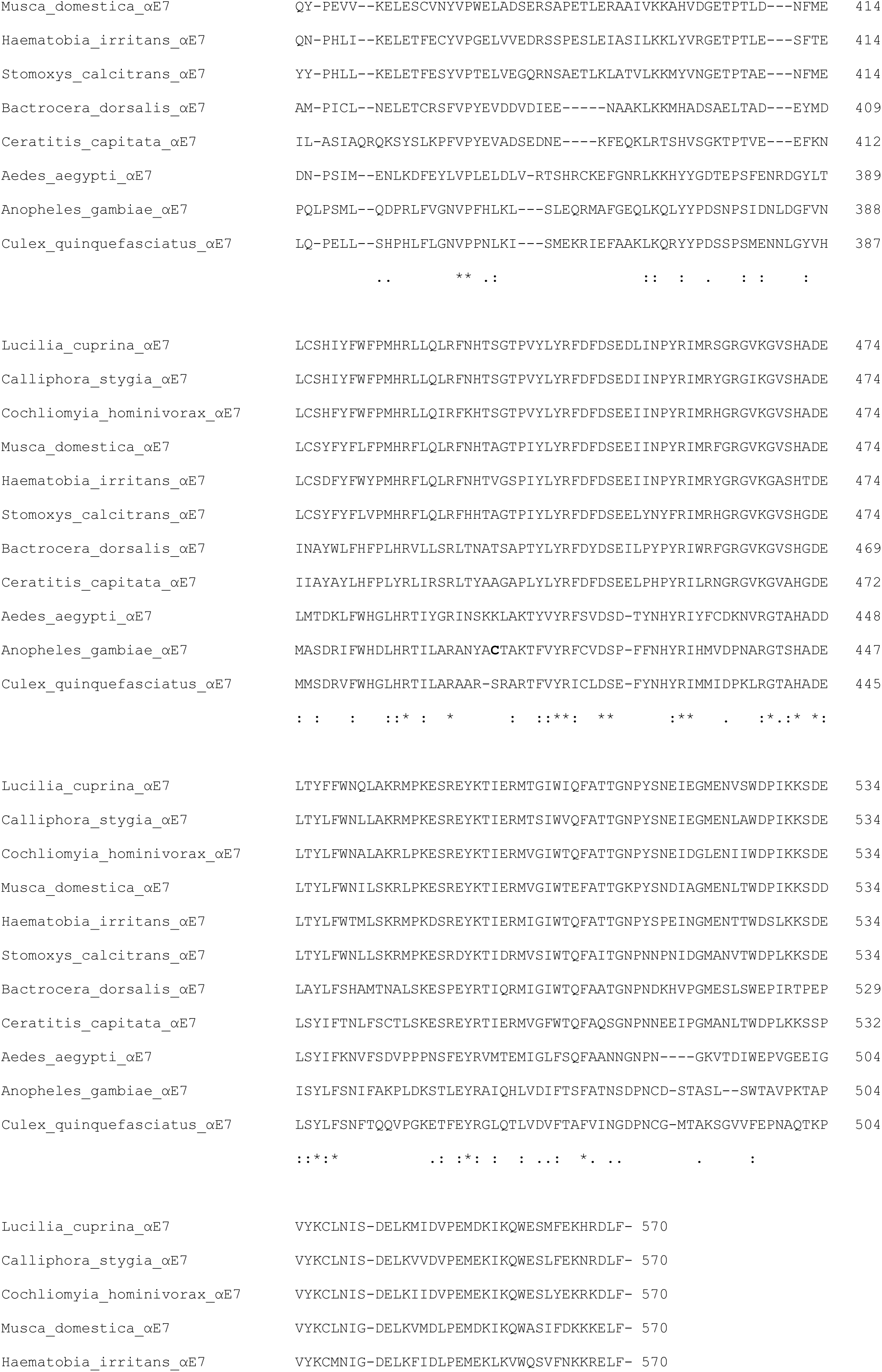
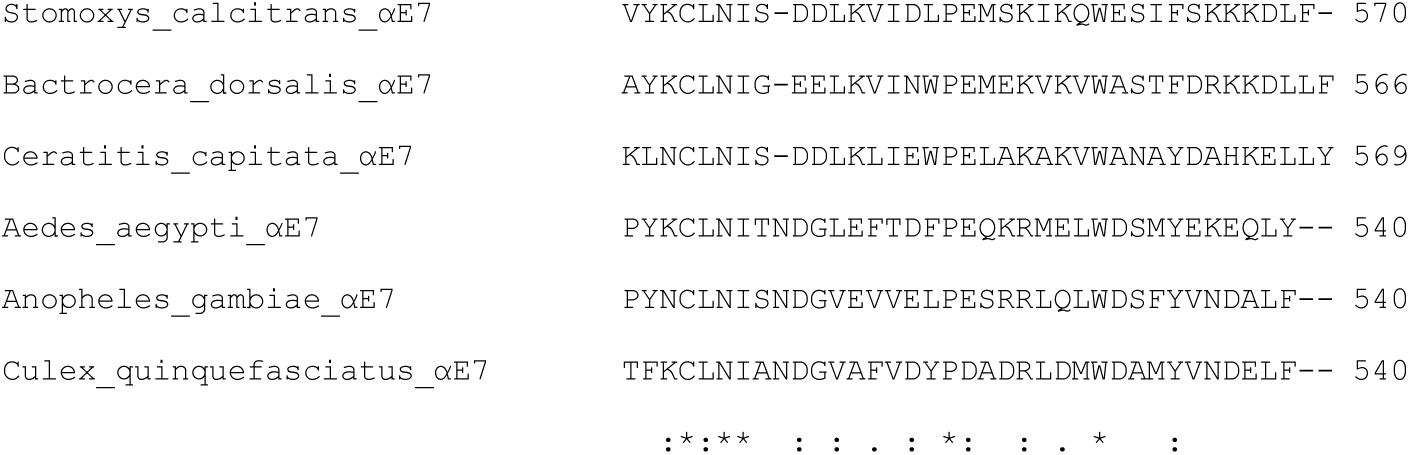
Multiple sequence alignment of carboxylic ester hydrolases from insect species.

**Supplementary figure 3:**
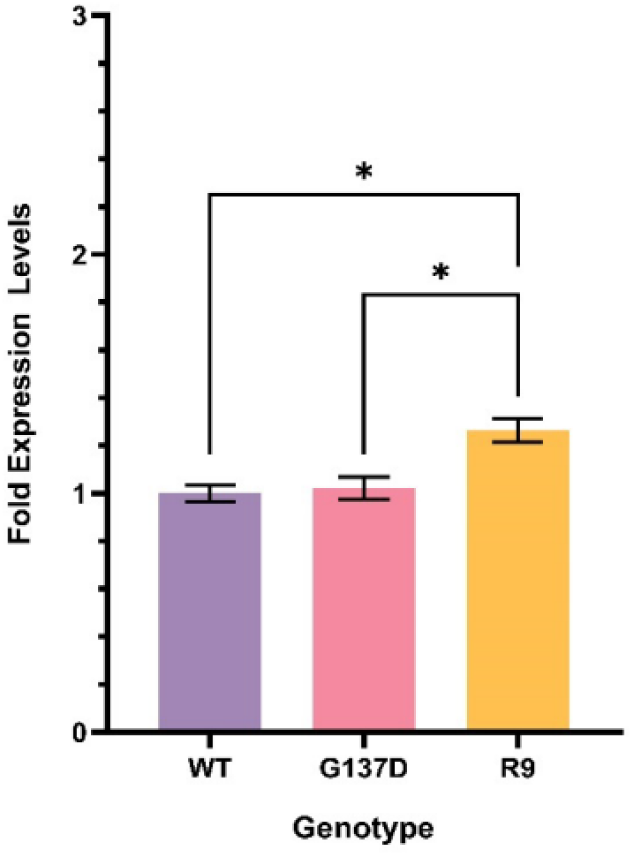
Relative expression levels of *Lcα*E7 transgenes. The expression of the *Lcα*E7 genes were normalised to the expression of housekeeping genes *Rpl11* and *CG13220*. The fold expression levels shown are relative to those measured for the WT gene. No expression of *Lcα*E7 was detected in the control line, hence the fold expression level cannot be calculated. The data represented: mean ± SEM; (n=3); Tukey’s HSD test (P≤0.05 (*)).

**Supplementary table 1:**
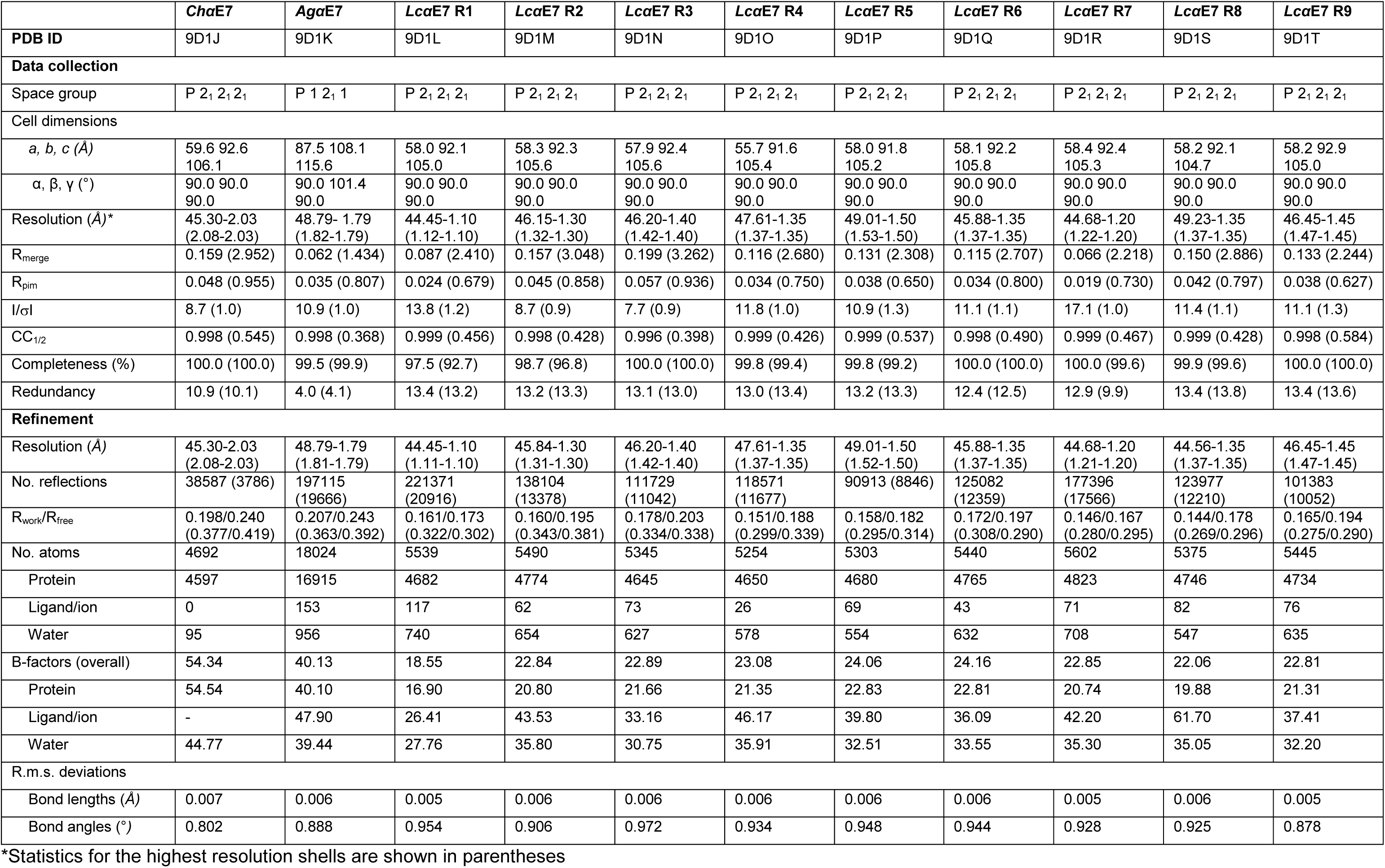
Crystallographic table of statistics.

**Supplementary table 2:**
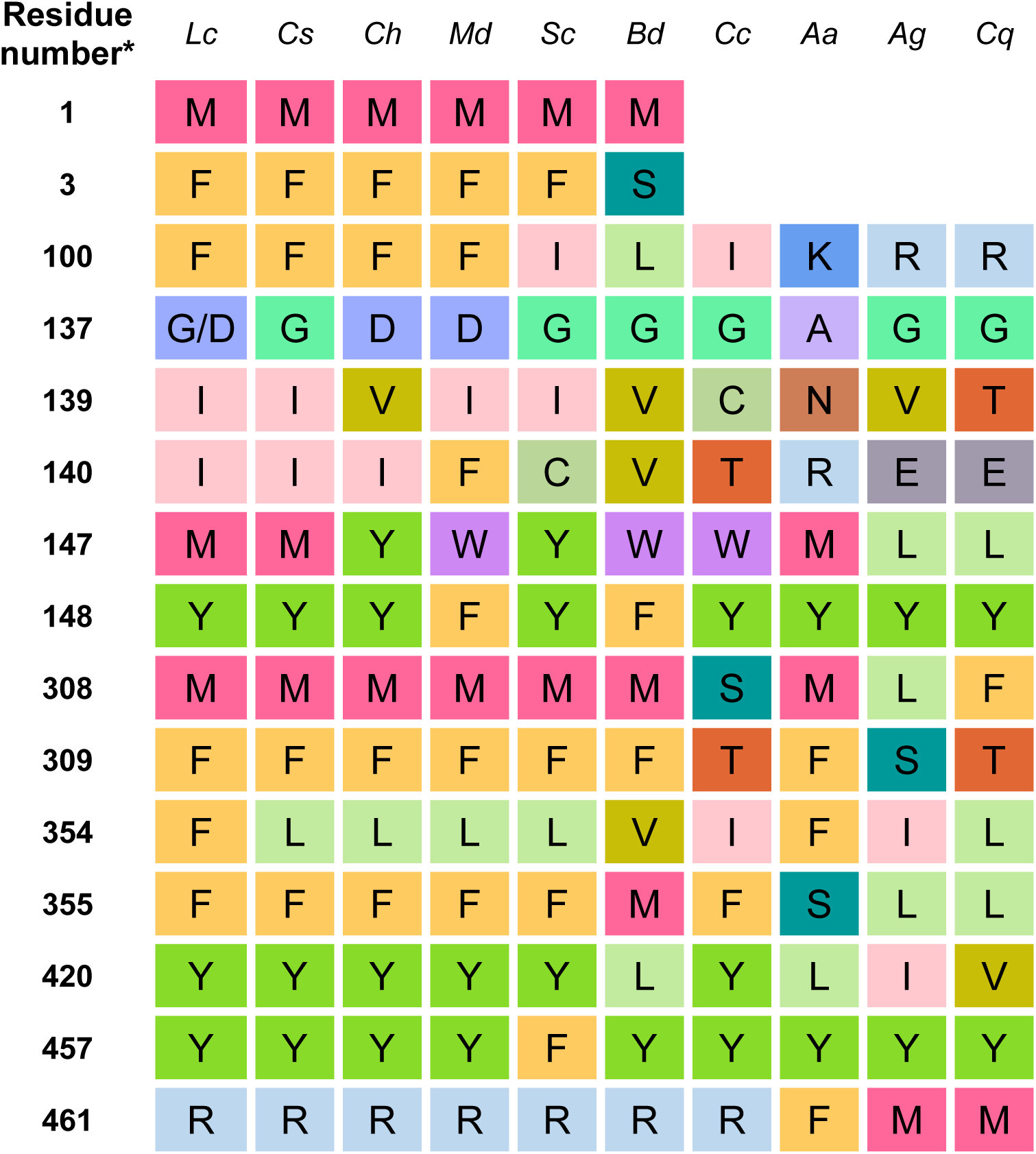
Comparison of active site residues in orthologs. Cells are coloured by amino acid for ease of comparison. *or equivalent residue.

**Supplementary table 3:**
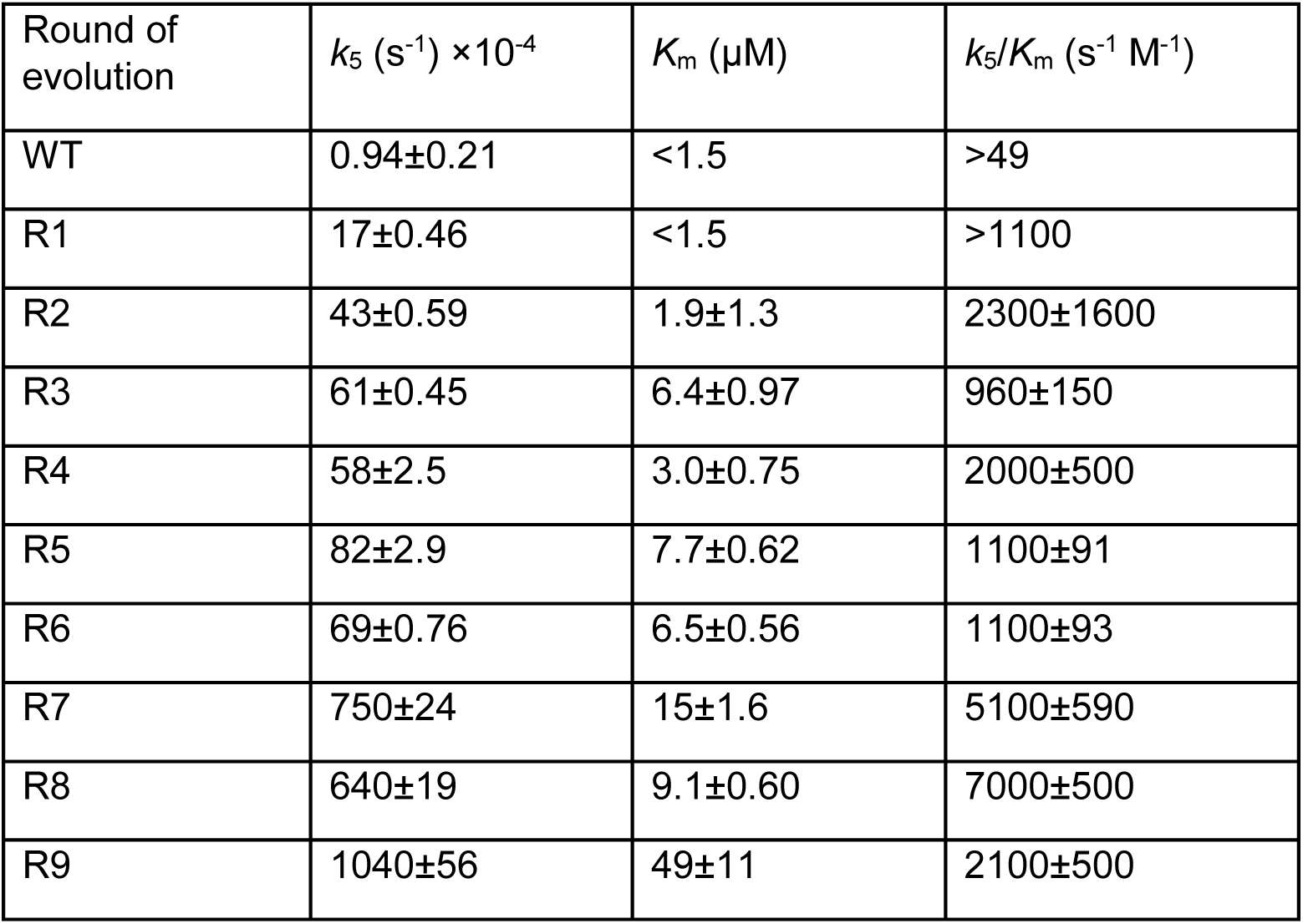
Michaelis-Menten kinetic parameters for hydrolysis of diethyl umbelliferyl phosphate (organophosphate). Values are mean and standard error (n=4).

**Supplementary table 4:**
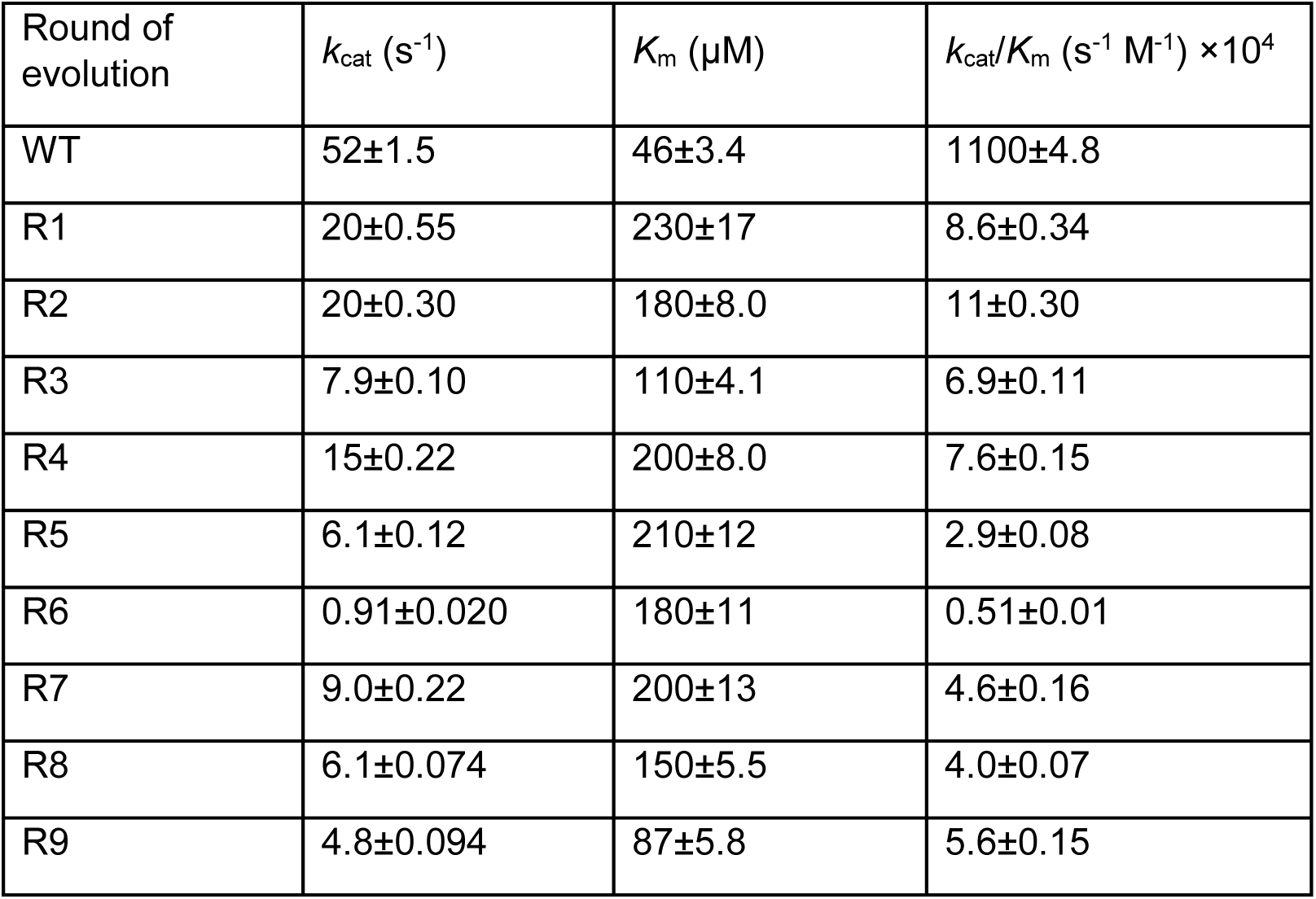
Michaelis-Menten kinetic parameters for hydrolysis of 4-nitrophenyl butyrate (carboxylester). Values are mean ± standard error (n=4)

**Supplementary table 5:**
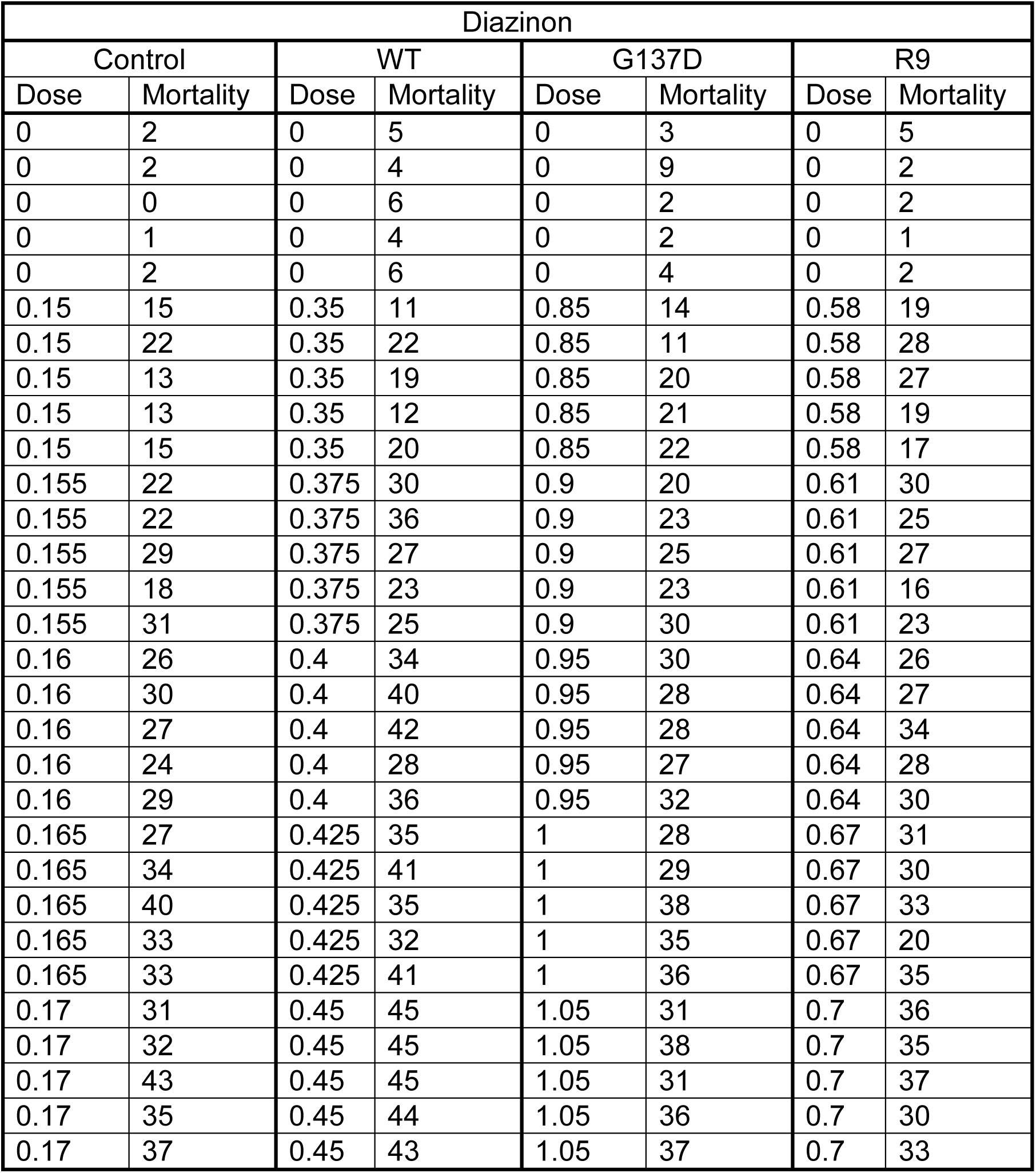
Toxicology data of Diazinon on *Drosophila melanogaster.* Sample size =50.

**Supplementary Table 6:**
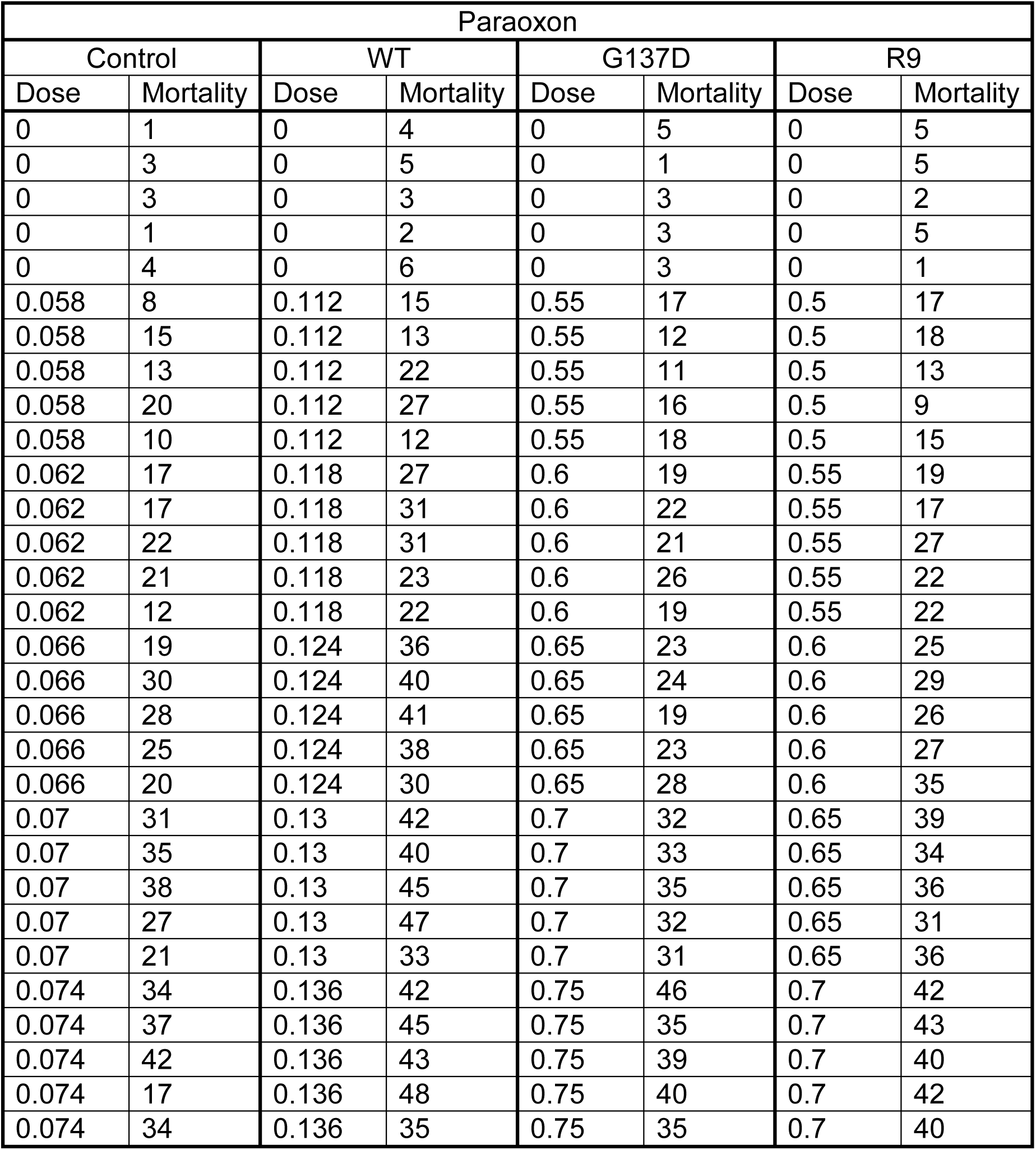
Toxicology data of Paraoxon on *Drosophila melanogaster.* Sample size =50.

**Supplementary table 7:**
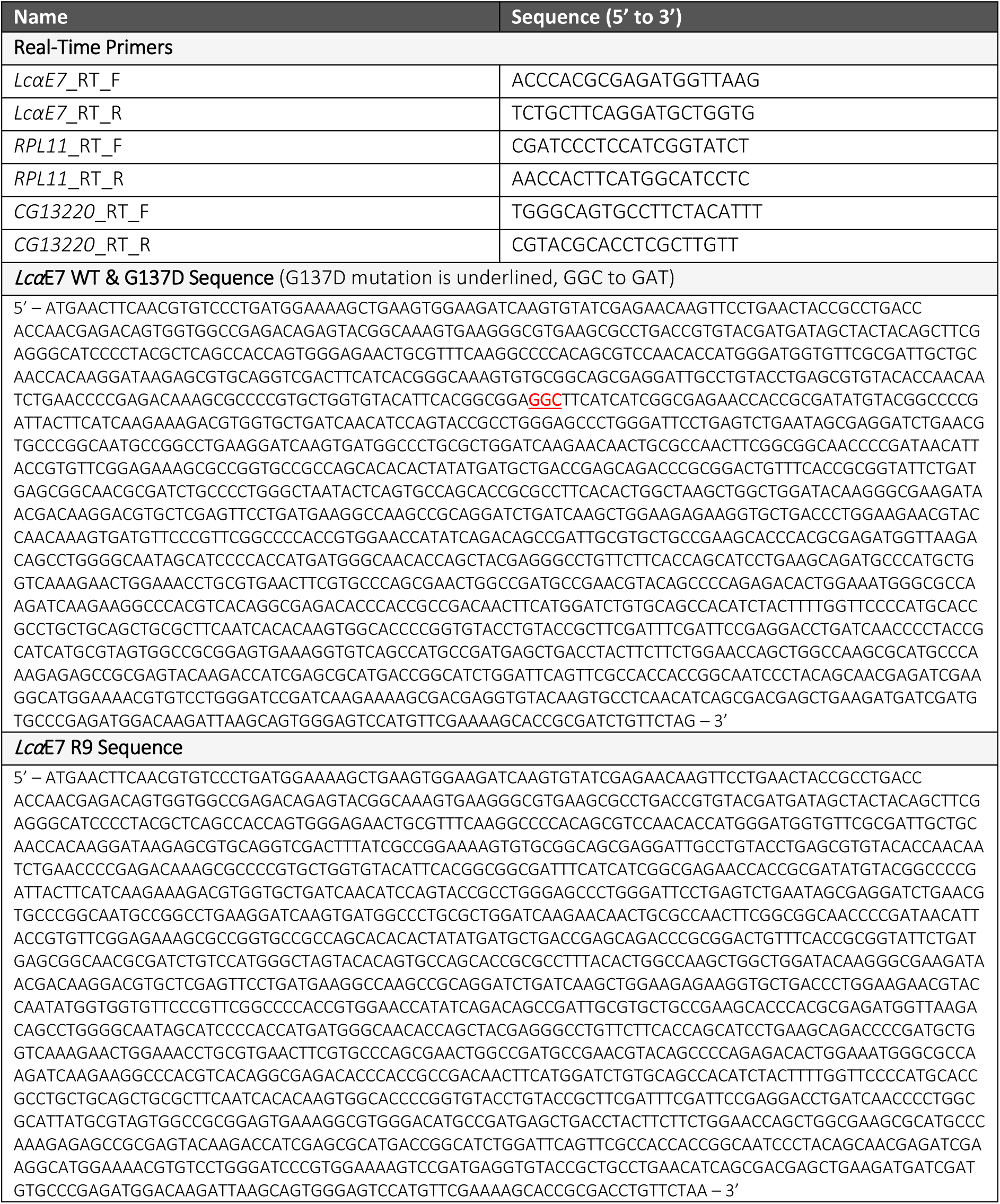
Primers and *LcαE7* sequences used.

**Supplementary table 8:** List of *Drosophila melanogaster lines*.

| Lines | Description |
| --- | --- |
| 25709w | Contains 2nd Chromosome attP40 docking site for phiC31 integrase-mediated transformation (Bloomington Stock #25709) and an X-chromosome with a *white* mutation and phiC31 construct (Bloomington Stock #24749). |
| HR-GAL4 | Contains *Accord-GAL4* on 3rd Chromosome. |
| **Transgenic Injected Lines (UAS- *LcaE7*)** |  |
| *UAS-Control* | Control line with *UAS-Null* on 2nd Chromosome. *UAS-Null* is a construct with no inserted gene. |
| *UAS-WT* | Line with *LcaE7 UAS-WT* on 2nd Chromosome. |
| *UAS-G137D* | Line with *LcaE7 UAS-G137D* on 2nd Chromosome. |
| *UAS-R9* | Line with *LcaE7 UAS-R9* on 2nd Chromosome. |
| **Transgenic Overexpression Lines** |  |
| Control | Cross of *UAS-25709* with *HR-GAL4* lines. Control line with same genetic background as the overexpression lines. |
| WT | Cross of *UAS-WT* with *HR-GAL4* lines. *LcaE7* WT overexpression under the control of *Accord*. |
| G137D | Cross of *UAS-G137D* with *HR-GAL4* lines. *LcaE7* G137D overexpression under the control of *Accord*. |
| R9 | Cross of *UAS-R9* with *HR-GAL4* lines. *LcaE7* R9 overexpression under the control of *Accord*. |

## References

1. Y. H. Kim, S. H. Lee, Which acetylcholinesterase functions as the main catalytic enzyme in the Class Insecta? Insect Biochem Mol Biol 43, 47–53 (2013).

2. J. G. Oakeshott et al., How many genetic options for evolving insecticide resistance in heliothine and spodopteran pests? Pest Manag Sci 69, 889–896 (2013).

3. J. Liang et al., Insect Resistance to Insecticides: Causes, Mechanisms, and Exploring Potential Solutions. Arch Insect Biochem Physiol 118, e70045 (2025).

4. R. Feyereisen, W. Dermauw, T. Van Leeuwen, Genotype to phenotype, the molecular and physiological dimensions of resistance in arthropods. Pestic Biochem Physiol 121, 61–77 (2015).

5. R. D. Newcomb et al., A single amino acid substitution converts a carboxylesterase to an organophosphorus hydrolase and confers insecticide resistance on a blowfly. Proc Natl Acad Sci U S A 94, 7464–7468 (1997).

6. L. W. Bergamo, P. Fresia, A. M. Azeredo-Espin, Incongruent nuclear and mitochondrial genetic structure of new world screwworm fly populations due to positive selection of mutations associated with dimethyl- and diethyl-organophosphates resistance. PLoS One 10, e0128441 (2015).

7. C. Claudianos, R. J. Russell, J. G. Oakeshott, The same amino acid substitution in orthologous esterases confers organophosphate resistance on the house fly and a blowfly. Insect Biochemistry and Molecular Biology 29, 675–686 (1999).

8. R. A. de Carvalho, T. T. Torres, A. M. de Azeredo-Espin, A survey of mutations in the Cochliomyia hominivorax (Diptera: Calliphoridae) esterase E3 gene associated with organophosphate resistance and the molecular identification of mutant alleles. Vet Parasitol 140, 344–351 (2006).

9. R. Heidari et al., Hydrolysis of organophosphorus insecticides by in vitro modified carboxylesterase E3 from Lucilia cuprina. Insect Biochem Mol Biol 34, 353–363 (2004).

10. R. D. Newcomb, D. M. Gleeson, C. G. Yong, R. J. Russell, J. G. Oakeshott, Multiple mutations and gene duplications conferring organophosphorus insecticide resistance have been selected at the Rop-1 locus of the sheep blowfly, Lucilia cuprina. J Mol Evol 60, 207–220 (2005).

11. A. G. Davies et al., Scalloped wings is the Lucilia cuprina Notch homologue and a candidate for the modifier of fitness and asymmetry of diazinon resistance. Genetics 143, 1321–1337 (1996).

12. C. J. Hartley et al., Amplification of DNA from preserved specimens shows blowflies were preadapted for the rapid evolution of insecticide resistance. Proc Natl Acad Sci U S A 103, 8757–8762 (2006).

13. P. M. Campbell, R. D. Newcomb, R. J. Russell, J. G. Oakeshott, Two different amino acid substitutions in the ali-esterase, E3, confer alternative types of organophosphorus insecticide resistance in the sheep blowfly, Lucilia cuprina. Insect Biochemistry and Molecular Biology 28, 139–150 (1998).

14. R. Qu, J. Zhu, M. Li, R. Jashenko, X. Qiu, Multiple Genetic Mutations Related to Insecticide Resistance are Detected in Field Kazakhstani House Flies (Muscidae: Diptera). J Med Entomol 58, 2338–2348 (2021).

15. T. R. Fukuto, Mechanism of action of organophosphorus and carbamate insecticides. Environ Health Perspect 87, 245–254 (1990).

16. T. C. Marrs, Organophosphate poisoning. Pharmacol Ther 58, 51–66 (1993).

17. D. H. Hopkins et al., Structure of an Insecticide Sequestering Carboxylesterase from the Disease Vector Culex quinquefasciatus: What Makes an Enzyme a Good Insecticide Sponge? Biochemistry 56, 5512–5525 (2017).

18. L. Grigoraki et al., Functional and immunohistochemical characterization of CCEae3a, a carboxylesterase associated with temephos resistance in the major arbovirus vectors Aedes aegypti and Ae. albopictus. Insect Biochem Mol Biol 74, 61–67 (2016).

19. Y. H. Gong et al., Functional characterization of carboxylesterase gene mutations involved in Aphis gossypii resistance to organophosphate insecticides. Insect Mol Biol 26, 702–714 (2017).

20. F. D. Guerrero, Cloning of a horn fly cDNA, HialphaE7, encoding an esterase whose transcript concentration is elevated in diazinon-resistant flies. Insect Biochem Mol Biol 30, 1107–1115 (2000).

21. L. L. Wang, X. P. Lu, G. Smagghe, L. W. Meng, J. J. Wang, Functional characterization of BdB1, a well-conserved carboxylesterase among tephritid fruit flies associated with malathion resistance in Bactrocera dorsalis (Hendel). Comp Biochem Physiol C Toxicol Pharmacol 200, 1–8 (2017).

22. P. D. Mabbitt et al., Conformational Disorganization within the Active Site of a Recently Evolved Organophosphate Hydrolase Limits Its Catalytic Efficiency. Biochemistry 55, 1408–1417 (2016).

23. C. J. Jackson et al., Structure and function of an insect alpha-carboxylesterase (alphaEsterase7) associated with insecticide resistance. Proc Natl Acad Sci U S A 110, 10177–10182 (2013).

24. G. J. Correy et al., Mapping the Accessible Conformational Landscape of an Insect Carboxylesterase Using Conformational Ensemble Analysis and Kinetic Crystallography. Structure 24, 977–987 (2016).

25. G. J. Correy et al., Overcoming insecticide resistance through computational inhibitor design. Proc Natl Acad Sci U S A 116, 21012–21021 (2019).

26. A. Couce et al., Changing fitness effects of mutations through long-term bacterial evolution. Science 383, eadd1417 (2024).

27. M. L. M. Salverda et al., Initial Mutations Direct Alternative Pathways of Protein Evolution. PLOS Genetics 7, e1001321 (2011).

28. D. Aggeli, Y. Li, G. Sherlock, Changes in the distribution of fitness effects and adaptive mutational spectra following a single first step towards adaptation. Nat Commun 12, 5193 (2021).

29. G. Yang et al., Higher-order epistasis shapes the fitness landscape of a xenobiotic-degrading enzyme. Nat Chem Biol 15, 1120–1128 (2019).

30. T. N. Starr, L. K. Picton, J. W. Thornton, Alternative evolutionary histories in the sequence space of an ancient protein. Nature 549, 409–413 (2017).

31. C. M. Miton, N. Tokuriki, How mutational epistasis impairs predictability in protein evolution and design. Protein Sci 25, 1260–1272 (2016).

32. F. Cui et al., Carboxylesterase-mediated insecticide resistance: Quantitative increase induces broader metabolic resistance than qualitative change. Pestic Biochem Physiol 121, 88–96 (2015).

33. D. M. Weinreich, N. F. Delaney, M. A. DePristo, D. L. Hartl, Darwinian Evolution Can Follow Only Very Few Mutational Paths to Fitter Proteins. Science 312, 111–114 (2006).

34. N. Tokuriki et al., Diminishing returns and tradeoffs constrain the laboratory optimization of an enzyme. Nature Communications 3, 1257 (2012).

35. C. M. Miton et al., Evolutionary repurposing of a sulfatase: A new Michaelis complex leads to efficient transition state charge offset. Proceedings of the National Academy of Sciences 115, E7293–E7302 (2018).

36. E. Campbell et al., The role of protein dynamics in the evolution of new enzyme function. Nature Chemical Biology 12, 944–950 (2016).

37. N. J. Hawkins, C. Bass, A. Dixon, P. Neve, The evolutionary origins of pesticide resistance. Biological Reviews 94, 135–155 (2019).

38. P. J. Wijngaarden, F. van den Bosch, M. J. Jeger, R. F. Hoekstra, Adaptation to the cost of resistance: a model of compensation, recombination, and selection in a haploid organism. Proc Biol Sci 272, 85–89 (2005).

39. H. Chung et al., Cis-regulatory elements in the Accord retrotransposon result in tissue-specific expression of the Drosophila melanogaster insecticide resistance gene Cyp6g1. Genetics 175, 1071–1077 (2007).

40. Y. Boublik et al., Acetylcholinesterase engineering for detection of insecticide residues. Protein Engineering 15, 43–50 (2002).

41. J. Vontas et al., Insecticide resistance in Tephritid flies. Pesticide Biochemistry and Physiology 100, 199–205 (2011).

42. J. A. McKenzie, G. M. Clarke, Diazinon resistance, fluctuating asymmetry and fitness in the Australian sheep blowfly, lucilia cuprina. Genetics 120, 213–220 (1988).

43. Z. Chen, R. Newcomb, E. Forbes, J. McKenzie, P. Batterham, The acetylcholinesterase gene and organophosphorus resistance in the Australian sheep blowfly, Lucilia cuprina. Insect Biochemistry and Molecular Biology 31, 805–816 (2001).

44. R. V. Rane et al., Detoxifying enzyme complements and host use phenotypes in 160 insect species. Current opinion in insect science 31, 131–138 (2019).

45. H. Chung et al., Characterization of Drosophila melanogaster cytochrome P450 genes. Proceedings of the National Academy of Sciences 106, 5731–5736 (2009).

46. I. Rayment, "[12] Reductive alkylation of lysine residues to alter crystallization properties of proteins" in Methods in Enzymology. (Academic Press, 1997), vol. 276, pp. 171–179.

47. D. A.-O. Aragão et al., MX2: a high-flux undulator microfocus beamline serving both the chemical and macromolecular crystallography communities at the Australian Synchrotron. Journal of synchrotron radiation 25, 885–891 (2018).

48. W. Kabsch, XDS. Acta Crystallographica Section D: Biological Crystallography 66, 125–132 (2010).

49. P. R. Evans, G. N. Murshudov, How good are my data and what is the resolution? Crystallogr D Biol Crystallogr 69, 1204–1214 (2013).

50. A. J. McCoy et al., Phaser crystallographic software. J Appl Crystallogr 40, 658–674 (2007).

51. P. H. Zwart et al., Automated structure solution with the PHENIX suite. Methods in Molecular Biology 426, 419–435 (2008).

52. P. Emsley, K. Cowtan, Coot: model-building tools for molecular graphics. Acta Crystallogr D Biol Crystallogr 60, 2126–2132 (2004).

53. C. J. Williams et al., MolProbity: More and better reference data for improved all-atom structure validation. Protein Science 27, 293–315 (2018).

54. J. Bischof, R. K. Maeda, M. Hediger, F. Karch, K. Basler, An optimized transgenesis system for Drosophila using germ-line-specific C31 integrases. Proceedings of the National Academy of Sciences 104, 3312–3317 (2007).

55. A. H. Brand, N. Perrimon, Targeted gene expression as a means of altering cell fates and generating dominant phenotypes. Development 118, 401–415 (1993).

56. T. Perry et al., Effects of mutations in Drosophila nicotinic acetylcholine receptor subunits on sensitivity to insecticides targeting nicotinic acetylcholine receptors. Pesticide Biochemistry and Physiology 102, 56–60 (2012).

57. W. S. Abbott, A Method of Computing the Effectiveness of an Insecticide. Journal of Economic Entomology 18, 265–267 (1925).

58. D. J. Finney, Probit analysis: A statistical treatment of the sigmoid response curve, 2nd ed (Cambridge University Press, New York, NY, US, 1952), pp. xiv, 318-xiv, 318.

