## Supplemental Information for "Structural and Evolutionary Constraints of Organophosphate Resistance in Dipteran Carboxylesterases"

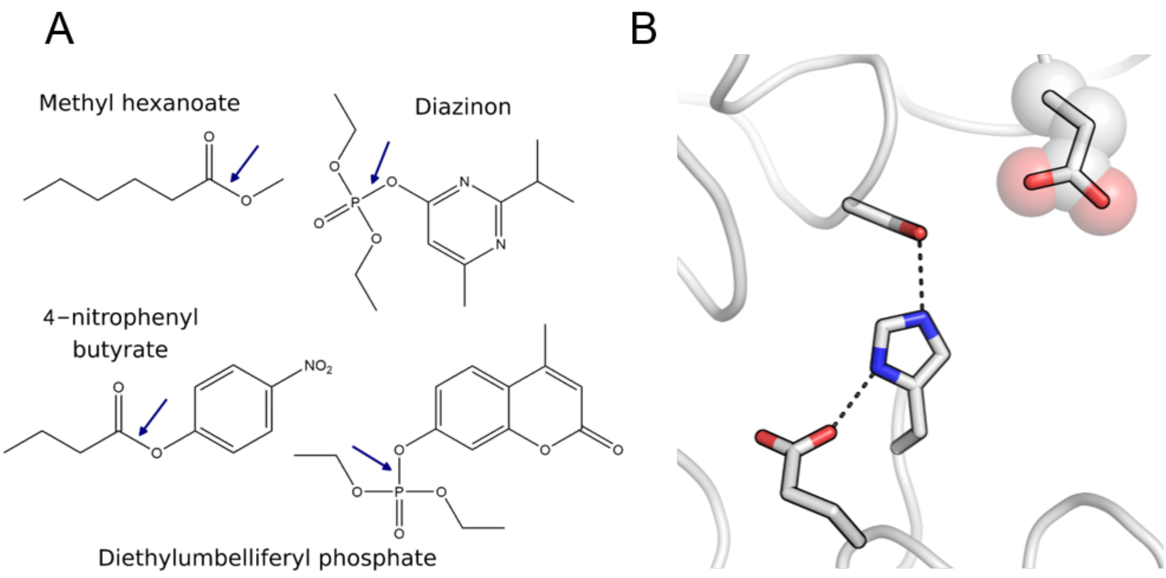


**Supplementary figure 1: (A)** a substrate of *L. cuprina αE7* (Methyl hexanoate), the diethyl-OP insecticide (Diazinon), the chromogenic ester-substrate (4-nitrophenbyl butyrate) and the fluorogenic OP-substrate (Diethylumbelliferyl phosphate). **(B)** Catalytic triad (Ser218, Glu351 His471) and Asp137 in *Lcα*E7 (PDB 5C8V).

```
Lucilia_cuprina_αE7               MNFNVSLMEKLKWKIKCIENKFLNYRLTTNETVVAETEYGKVKGVKRLTVYDDSYYSFEG   60
Calliphora_stygia_αE7             MNFNASLMEKLRWKIKCIEYKFQNYRLTTNETVVAETEYGKVKGIKRLTIYDDSYYSFEG   60
Cochliomyia_hominivorax_αE7       MNFKVSQVEKLKWKIKCFENKFLNYRLSTNETAVAETEYGKVKGIKRLTVYDDSYYSFEG   60
Musca_domestica_αE7               MNFKVSQMERLSWKLKCMVNKYTNYRLSTNETQIIDTEYGQIKGVKRMTVYDDSYYSFES   60
Haematobia_irritans_αE7           MNFNVSFLEKLRWKIKCVENKILNYRLITNETHIVDTEYGKIKGVKRLTVYDDSFYSFEG   60
Stomoxys_calcitrans_αE7           MNFNNSFLERLRWKIKCIENKLTTYRLVTNETQIVDTEYGKIKGVKRLNVYDESFYSFEG   60
Bactrocera_dorsalis_αE7           MHSNVGFLDKLRWRLNVLGNRYAQYRLTTNETAVVDTEYGKVKGVKRRTIYEVPYYSFEG   60
Ceratitis_capitata_αE7            MQSNIGFIEKFRWRLKVYEHKYQQNRLATAETLIVETEYGKVEGIKRLSIYNIPYYSFEG   60
Aedes_aegypti_αE7                 ---------------------------MSAESPVVCTKYGPVKGIRKTAATGVEYFSFQR   33
Anopheles_gambiae_αE7             ---------------------------MANSELIVSTGYGPVQGTARTSLYGTGYVSFQG   33
Culex_quinquefasciatus_αE7        ---------------------------MSLESLTVQTKYGPVRGKRSVSLLGQEYVSFQG   33
                                                              : .     * ** :.*          : **:

Lucilia_cuprina_αE7               IPYAQPPVGELRFKAPQRPTPWDGVRDCCNHKDKSVQVDFITGKVCGSEDCLYLSVYTNN   120
Calliphora_stygia_αE7             IPYAQPPLGELRFKAPQRPTPWDGERDCCNNRDKSVQRDFITGKVCGSEDCLYLSVYTNN   120
Cochliomyia_hominivorax_αE7       IPYAQPPLGELRFKAPQRPTPWDGVRDCCNNKDKSVQVDFITGKTCGSEDCLYLSVYTNN   120
Musca_domestica_αE7               IPYAKPPVGELRFKAPQRPVPWEGVRDCCGPANRSVQTDFISGKPTGSEDCLYLNVYTND   120
Haematobia_irritans_αE7           IPYAKPPVGELRFKAPQRPVPWDGVKDCCHAASRSVQTDFISGNSSGSEDCLYLNVYTNN   120
Stomoxys_calcitrans_αE7           IPYAQPPLGELRFKAPQRPVPWEGVKDCVTAAPKSVQTDIISGKSTGSEDCLYLNVYTNN   120
Bactrocera_dorsalis_αE7           IPYAKPPVDELRFKAPERPEPWDGVLDCLSPKDRAVQLHLITNNAEGSEDCLYLNVYAKN   120
Ceratitis_capitata_αE7            IPYAQPPVGELRFRAPQRPTPWEGVRDCKSTKEMAVQTHIITGILEGSEDCLYLNVYTNN   120
Aedes_aegypti_αE7                 IPYAKPPVGDLRFKDAVPPAAWTEELDCTVQGPAGYQFSKLQNKIIGNEDCLHMNVFTKS   93
Anopheles_gambiae_αE7             IPYAKPPVGELRFKDPTPPENWTQVLDCTEQCDPCFHFDRRVNKIVGSEDSLRLNIFSKT   93
Culex_quinquefasciatus_αE7        IPYARAPEGELRFKAPVPPQNWTETLDCSQQCEPCYHFDRRLQKIVGCEDSLKINVFAKE   93
                                  ****: * .:***:    *  *    **        :         * **.* :.::::

Lucilia_cuprina_αE7               LNPETKRPVLVYIHGGGFIIGENHRDMYGPDYFIKKDVVLINIQYRLGALGFLSLNSEDL   180
Calliphora_stygia_αE7             LTPETKRPVLVYIHGGGFIIGENHRDMYGPDYLMKKDVVVINIQYRLGVLGFLSLNTEDL   180
Cochliomyia_hominivorax_αE7       LAPETKRPVLVYIHGGDFVIGENHREYYGPDYFIKKDVVLITIQYRLGVLGFLSLNSEEL   180
Musca_domestica_αE7               LNPDKKRPVMVFIHGGDFIFGEANRNWFGPDYFMKKPVVLVTVQYRLGVLGFLSLKSENL   180
Haematobia_irritans_αE7           LNTDTKRPVLVFFHGGGFICGEANRNYYGADYFIKKDVVFITVQYRLGVLGFLSLNSENL   180
Stomoxys_calcitrans_αE7           LNPEKKRPVMVFLHGGGFICGEANENYYGGDYFIKKDVILVTVQYRLGILGFLSLKSEDL   180
Bactrocera_dorsalis_αE7           LKPDKPQPVLVWIHGGGFVVGEANRDWFGPDYFMEKDVVLVTVQYRLGVFGFLTLTSPEL   180
Ceratitis_capitata_αE7            TLPDKPRPVMIWIHGGGLCTGEATREWYGPDYFMQKDIVLVTMQYRLGVLGFLSLGTPEL   180
Aedes_aegypti_αE7                 LDKGERLPVMLYIHGGAFNRGSSGVEMYGPDYLIQADVVFVSFNYRIGALGFISFESPEV   153
Anopheles_gambiae_αE7             IKPTKPLPVMVYIYGGGFVEGTSGTELYGPDYLIEKDIVLVTLNYRVGALGFLCCQSPTA   153
Culex_quinquefasciatus_αE7        INPSKPLPVMLYIYGGGFTEGTSGTELYGPDFLVQKDIVLVSFNYRIGALGFLCCQSEQD   153
                                         **:::::** :  *    : :* *::::  ::.:..:**:* :**:   :

Lucilia_cuprina_αE7               NVPGNAGLKDQVMALRWIKNNCANFGGNPDNITVFGESAGAASTHYMMLTEQTRGLFHRG   240
```

```
Calliphora_stygia_αE7              NVPGNAGLKDQVMALRWIKNNCANFGGNPDNITVFGESAGGASTHYMMLTEQTRGLFHRG   240
Cochliomyia_hominivorax_αE7        NVPGNAGLKDQVMALRWIKNNCANFGGNPDNITVFGESAGGASAHYMMLTEQTRGLFHRG   240
Musca_domestica_αE7                NVPGNAGLKDQVMALRWVKSNIANFGGDVDNITVFGESAGGASTHYMMITEQTRGLFHRG   240
Haematobia_irritans_αE7            NVPGNAGLKDQVMALRWIKNNCASFGGDPDCITLFGESAGAASTHYMMITEQARGLFHRA   240
Stomoxys_calcitrans_αE7            NIPGNAALKDIVMALRWVKNNCTNFGGDPDNITLFGESAGSASTHYMMITEQTRGLFHRA   240
Bactrocera_dorsalis_αE7            NIPGNAGLKDQVLALKWVKNNIANFGGDPNCITVFGESAGAASTHYLSITEQTRGLFHRA   240
Ceratitis_capitata_αE7             NVPGNSGLKDQVLAIKWVKNNCARFGGNPDCITVFGESAGATSAHCMMLTEQTQGLFHRA   240
Aedes_aegypti_αE7                  DLPGNAGLKDQNLALRWVVENIEAFGGDPNNITLFGESAGGCSVHYHMISDQSKGLFQRA   213
Anopheles_gambiae_αE7              GVPGNAGLKDQRLALRWVRDNIASFGGDPSAITLFGHSAGGASVQYHTIADASKNLFQRA   213
Culex_quinquefasciatus_αE7         GVPGNAGLKDQNLAIRWVLENIAAFGGDPKRVTLVGHSAGAASVQYHLISDASKDLFQRA   213
                                   .:***:.***  :*::*: .*   ***: . :*:.*.***. *.:   ::: ::.**:*.

Lucilia_cuprina_αE7                ILMSGNAICPWANTQCQHRAFTLAKLAGYKGEDNDKDVLEFLMKAKPQDLIKLEEKVLTL   300
Calliphora_stygia_αE7              VLMSGNAICPWASTECQHRAYNIAKLAGYKGENNDKDVLEFLLKVKSQDLIKLEDKVLTP   300
Cochliomyia_hominivorax_αE7        ILMSGNAVCPWAISQNQHRAYAIAKLNGYKGENNDKDVLEFLMKAKAHDLIKLEDKVLTP   300
Musca_domestica_αE7                IMMSGNSMCSWASTECQSRALTMAKRVGYKGEDNEKDILEFLMKANPYDLIKEEPQVLTP   300
Haematobia_irritans_αE7            VLMSGTAMCIWAHTQCQHRGYTIAKRIGYKGENNDKDVYDFLMKANPYDLAREEHKVLTN   300
Stomoxys_calcitrans_αE7            ILMSGTAMCIWAHTECQHRAFAIAKRLGYKGEDNDKDVLEFLMKAHPYQMAKEEHLVLTP   300
Bactrocera_dorsalis_αE7            ILMSGAAIASWAHNGQERHAYPLAKLAGFKGEHDEQHVLEYLKKCKATDLAQLEQKVLTT   300
Ceratitis_capitata_αE7             ILMSGTALPLWETEDQKYRAFDLAKLAGYKGVDNDKDVLAYLRKCKAKDLIALEGRTLTA   300
Aedes_aegypti_αE7                  IVMSGCSLNNWSTIPRRQFSQRLAKALGWNGQGGDKAALEVLMKATPEDIVEKQGALRTE   273
Anopheles_gambiae_αE7              IIMSGSTMCSWALTPQRNWPEKLAKAIGWQGEGDEEAALQYLRQASPESIVDHQEKLFGP   273
Culex_quinquefasciatus_αE7         IVMSGSTYNSWSLTRQRNWVEKLAKAIGWDGQGGESGALRFLKAAKPEDIVANQEKLLTD   273
                                   ::*** :   *     .     :**  *:.*  .:.     *      .:   :

Lucilia_cuprina_αE7                EERTNKVMFPFGPTVEPYQTADCVLPKHPREMVKTAWGNSIPTMMGNTSYEGLFFTSILK   360
Calliphora_stygia_αE7              EEHKNKVMFAFGPTIEPYQSAECVIPKHPREMVKTAWGNSIPTMMGNTSYEGLLFYSIIK   360
Cochliomyia_hominivorax_αE7        EEHVNKVMFAFGPTVEPYQTADCVLPKHPREMVKTAWGNSIPTMMGNTSYEGLLFTPIVK   360
Musca_domestica_αE7                EEMQNKVMFPFGPTVEPYQTADCVVPKPIREMVKSAWGNSIPTLIGNTSYEGLLFKSIAK   360
Haematobia_irritans_αE7            EELRDKVMFAFGPTTEPYETPDCVLPKPNREMLKTAWGNSIPTLIGNTSYEGLLFISVGK   360
Stomoxys_calcitrans_αE7            EELADKKMFAFGPTTEPYQTADCVLPKAPREMVKTAWGNSIPTMIGNTSYEGLLFLPFPK   360
Bactrocera_dorsalis_αE7            VELQRKIMFPFAPCIEPYDTPDCVISKSPRDLMRTAWSNSMPILAGHTSAEGLVMLPFIK   360
Ceratitis_capitata_αE7             EDRARNISTPFVYCVEPYVTPECVIQKPIREMMRTAWGNAIPLLVGHASDEGLIFLQGAK   360
Aedes_aegypti_αE7                  NEKENHIMFEFGPVVEPYIKDNCIIPEDPLKMCRKAWSHDIDILIGGNSEEGLFSLSEIK   333
Anopheles_gambiae_αE7              QEIQEGLLSPFAPTIEPYESEVCFIPRSPFEMSRTAWGNSIDIMIGGTSEEGLILLPKVK   333
Culex_quinquefasciatus_αE7         QDMQDDIFTPFGPTVEPYLTEQCMIPKEPFEMARTAWGDKIDIMIGGTSEEGLLLLQKIK   333
                                    :        *    *** .  *.: .   .: :.**.. :  : *  * ***.     *

Lucilia_cuprina_αE7                QM-PMLV--KELETCVNFVPSELADAERTAPETLEMGAKIKKAHVTGETPTAD---NFMD   414
Calliphora_stygia_αE7              QM-PAVL--KEMDTCANFVPPELADAERTAPETLEMGAKIKKAHTTGETPTAD---NFMD   414
Cochliomyia_hominivorax_αE7        QM-PALL--KELETCANFVPTELADSERSSAETLELGAKIKKAHVTGETPTND---NFLD   414
```

```
Musca_domestica_αE7               QY-PEVV--KELESCVNYVPWELADSERSAPETLERAAIVKKAHVDGETPTLD---NFME   414
Haematobia_irritans_αE7           QN-PHLI--KELETFECYVPGELVVEDRSSPESLEIASILKKLYVRGETPTLE---SFTE   414
Stomoxys_calcitrans_αE7           YY-PHLL--KELETFESYVPTELVEGQRNSAETLKLATVLKKMYVNGETPTAE---NFME   414
Bactrocera_dorsalis_αE7           AM-PICL--NELETCRSFVPYEVDDVDIEE-----NAAKLKKMHADSAELTAD---EYMD   409
Ceratitis_capitata_αE7            IL-ASIAQRQKSYSLKPFVPYEVADSEDNE----KFEQKLRTSHVSGKTPTVE---EFKN   412
Aedes_aegypti_αE7                 DN-PSIM--ENLKDFEYLVPLELDLV-RTSHRCKEFGNRLKKHYYGDTEPSFENRDGYLT   389
Anopheles_gambiae_αE7             PQLPSML--QDPRLFVGNVPFHLKL---SLEQRMAFGEQLKQLYYPDSNPSIDNLDGFVN   388
Culex_quinquefasciatus_αE7        LQ-PELL--SHPHLFLGNVPPNLKI---SMEKRIEFAAKLKQRYYPDSSPSMENNLGYVH   387
                                           ..       **  .:                ::  :  .   : :    :

Lucilia_cuprina_αE7               LCSHIYFWFPMHRLLQLRFNHTSGTPVYLYRFDFDSEDLINPYRIMRSGRGVKGVSHADE   474
Calliphora_stygia_αE7             LCSHIYFWFPMHRLLQLRFNHTSGTPVYLYRFDFDSEDIINPYRIMRYGRGIKGVSHADE   474
Cochliomyia_hominivorax_αE7       LCSHFYFWFPMHRLLQIRFKHTSGTPVYLYRFDFDSEEIINPYRIMRHGRGVKGVSHADE   474
Musca_domestica_αE7               LCSYFYFLFPMHRFLQLRFNHTAGTPIYLYRFDFDSEEIINPYRIMRFGRGVKGVSHADE   474
Haematobia_irritans_αE7           LCSDFYFWYPMHRFLQLRFNHTVGSPIYLYRFDFDSEEIINPYRIMRYGRGVKGASHTDE   474
Stomoxys_calcitrans_αE7           LCSYFYFLVPMHRFLQLRFHHTAGTPIYLYRFDFDSEELYNYFRIMRHGRGVKGVSHGDE   474
Bactrocera_dorsalis_αE7           INAYWLFHFPLHRVLLSRLTNATSAPTYLYRFDYDSEILPYPYRIWRFGRGVKGVSHGDE   469
Ceratitis_capitata_αE7            IIAYAYLHFPLYRLIRSRLTYAAGAPLYLYRFDFDSEELPHPYRILRNGRGVKGVAHGDE   472
Aedes_aegypti_αE7                 LMTDKLFWHGLHRTIYGRINSKKLAKTYVYRFSVDSD-TYNHYRIYFCDKNVRGTAHADD   448
Anopheles_gambiae_αE7             MASDRIFWHDLHRTILARANYACTAKTFVYRFCVDSP-FFNHYRIHMVDPNARGTSHADE   447
Culex_quinquefasciatus_αE7        MMSDRVFWHGLHRTILARAAR-SRARTFVYRICLDSE-FYNHYRIMMIDPKLRGTAHADE   445
                                  : :   :   ::* :  *      :  ::**:  **      :**   .   :*.:* *:

Lucilia_cuprina_αE7               LTYFFWNQLAKRMPKESREYKTIERMTGIWIQFATTGNPYSNEIEGMENVSWDPIKKSDE   534
Calliphora_stygia_αE7             LTYLFWNLLAKRMPKESREYKTIERMTSIWVQFATTGNPYSNEIEGMENLAWDPIKKSDE   534
Cochliomyia_hominivorax_αE7       LTYLFWNALAKRLPKESREYKTIERMVGIWTQFATTGNPYSNEIDGLENIIWDPIKKSDE   534
Musca_domestica_αE7               LTYLFWNILSKRLPKESREYKTIERMVGIWTEFATTGKPYSNDIAGMENLTWDPIKKSDD   534
Haematobia_irritans_αE7           LTYLFWTMLSKRMPKDSREYKTIERMIGIWTQFATTGNPYSPEINGMENTTWDSLKKSDE   534
Stomoxys_calcitrans_αE7           LTYLFWNLLSKRMPKESRDYKTIDRMVSIWTQFAITGNPNNPNIDGMANVTWDPLKKSDE   534
Bactrocera_dorsalis_αE7           LAYLFSHAMTNALSKESPEYRTIQRMIGIWTQFAATGNPNDKHVPGMESLSWEPIRTPEP   529
Ceratitis_capitata_αE7            LSYIFTNLFSCTLSKESREYRTIERMVGFWTQFAQSGNPNNEEIPGMANLTWDPLKKSSP   532
Aedes_aegypti_αE7                 LSYIFKNVFSDVPPPNSFEYRVMTEMIGLFSQFAANNGNPN----GKVTDIWEPVGEEIG   504
Anopheles_gambiae_αE7             ISYLFSNIFAKPLDKSTLEYRAIQHLVDIFTSFATNSDPNCD-STASL--SWTAVPKTAP   504
Culex_quinquefasciatus_αE7        LSYLFSNFTQQVPGKETFEYRGLQTLVDVFTAFVINGDPNCG-MTAKSGVVFEPNAQTKP   504
                                  ::*:*          .: :*: :  : ..:  *. ..        .     :

Lucilia_cuprina_αE7               VYKCLNIS-DELKMIDVPEMDKIKQWESMFEKHRDLF- 570
Calliphora_stygia_αE7             VYKCLNIS-DELKVVDVPEMEKIKQWESLFEKNRDLF- 570
Cochliomyia_hominivorax_αE7       VYKCLNIS-DELKIIDVPEMEKIKQWESLYEKRKDLF- 570
Musca_domestica_αE7               VYKCLNIG-DELKVMDLPEMDKIKQWASIFDKKKELF- 570
Haematobia_irritans_αE7           VYKCMNIG-DELKFIDLPEMEKLKVWQSVFNKKRELF- 570
```

```
Stomoxys_calcitrans_αE7          VYKCLNIS-DDLKVIDLPEMSKIKQWESIFSKKKDLF- 570

Bactrocera_dorsalis_αE7          AYKCLNIG-EELKVINWPEMEKVKVWASTFDRKKDLLF 566

Ceratitis_capitata_αE7           KLNCLNIS-DDLKLIEWPELAKAKVWANAYDAHKELLY 569

Aedes_aegypti_αE7                PYKCLNITNDGLEFTDFPEQKRMELWDSMYEKEQLY-- 540

Anopheles_gambiae_αE7            PYNCLNISNDGVEVVELPESRRLQLWDSFYVNDALF-- 540

Culex_quinquefasciatus_αE7       TFKCLNIANDGVAFVDYPDADRLDMWDAMYVNDELF-- 540

                                   :*:**  : : . : *:  : . *   :
```

**Supplementary figure 2:** Multiple sequence alignment of carboxylic ester hydrolases from insect species.

**Supplementary table 1:** Crystallographic table of statistics.

| | ***Chα*E7** | ***Agα*E7** | ***Lcα*E7 R1** | ***Lcα*E7 R2** | ***Lcα*E7 R3** | ***Lcα*E7 R4** | ***Lcα*E7 R5** | ***Lcα*E7 R6** | ***Lcα*E7 R7** | ***Lcα*E7 R8** | ***Lcα*E7 R9** |
|---|---|---|---|---|---|---|---|---|---|---|---|
| **PDB ID** | 9D1J | 9D1K | 9D1L | 9D1M | 9D1N | 9D1O | 9D1P | 9D1Q | 9D1R | 9D1S | 9D1T |
| **Data collection** | | | | | | | | | | | |
| Space group | P $2_1$ $2_1$ $2_1$ | P 1 $2_1$ 1 | P $2_1$ $2_1$ $2_1$ | P $2_1$ $2_1$ $2_1$ | P $2_1$ $2_1$ $2_1$ | P $2_1$ $2_1$ $2_1$ | P $2_1$ $2_1$ $2_1$ | P $2_1$ $2_1$ $2_1$ | P $2_1$ $2_1$ $2_1$ | P $2_1$ $2_1$ $2_1$ | P $2_1$ $2_1$ $2_1$ |
| Cell dimensions | | | | | | | | | | | |
| *a, b, c (Å)* | 59.6 92.6 106.1 | 87.5 108.1 115.6 | 58.0 92.1 105.0 | 58.3 92.3 105.6 | 57.9 92.4 105.6 | 55.7 91.6 105.4 | 58.0 91.8 105.2 | 58.1 92.2 105.8 | 58.4 92.4 105.3 | 58.2 92.1 104.7 | 58.2 92.9 105.0 |
| α, β, γ (°) | 90.0 90.0 90.0 | 90.0 101.4 90.0 | 90.0 90.0 90.0 | 90.0 90.0 90.0 | 90.0 90.0 90.0 | 90.0 90.0 90.0 | 90.0 90.0 90.0 | 90.0 90.0 90.0 | 90.0 90.0 90.0 | 90.0 90.0 90.0 | 90.0 90.0 90.0 |
| Resolution *(Å)** | 45.30-2.03 (2.08-2.03) | 48.79- 1.79 (1.82-1.79) | 44.45-1.10 (1.12-1.10) | 46.15-1.30 (1.32-1.30) | 46.20-1.40 (1.42-1.40) | 47.61-1.35 (1.37-1.35) | 49.01-1.50 (1.53-1.50) | 45.88-1.35 (1.37-1.35) | 44.68-1.20 (1.22-1.20) | 49.23-1.35 (1.37-1.35) | 46.45-1.45 (1.47-1.45) |
| $R_{merge}$ | 0.159 (2.952) | 0.062 (1.434) | 0.087 (2.410) | 0.157 (3.048) | 0.199 (3.262) | 0.116 (2.680) | 0.131 (2.308) | 0.115 (2.707) | 0.066 (2.218) | 0.150 (2.886) | 0.133 (2.244) |
| $R_{pim}$ | 0.048 (0.955) | 0.035 (0.807) | 0.024 (0.679) | 0.045 (0.858) | 0.057 (0.936) | 0.034 (0.750) | 0.038 (0.650) | 0.034 (0.800) | 0.019 (0.730) | 0.042 (0.797) | 0.038 (0.627) |
| $I/\sigma I$ | 8.7 (1.0) | 10.9 (1.0) | 13.8 (1.2) | 8.7 (0.9) | 7.7 (0.9) | 11.8 (1.0) | 10.9 (1.3) | 11.1 (1.1) | 17.1 (1.0) | 11.4 (1.1) | 11.1 (1.3) |
| $CC_{1/2}$ | 0.998 (0.545) | 0.998 (0.368) | 0.999 (0.456) | 0.998 (0.428) | 0.996 (0.398) | 0.999 (0.426) | 0.999 (0.537) | 0.998 (0.490) | 0.999 (0.467) | 0.999 (0.428) | 0.998 (0.584) |
| Completeness (%) | 100.0 (100.0) | 99.5 (99.9) | 97.5 (92.7) | 98.7 (96.8) | 100.0 (100.0) | 99.8 (99.4) | 99.8 (99.2) | 100.0 (100.0) | 100.0 (99.6) | 99.9 (99.6) | 100.0 (100.0) |
| Redundancy | 10.9 (10.1) | 4.0 (4.1) | 13.4 (13.2) | 13.2 (13.3) | 13.1 (13.0) | 13.0 (13.4) | 13.2 (13.3) | 12.4 (12.5) | 12.9 (9.9) | 13.4 (13.8) | 13.4 (13.6) |
| **Refinement** | | | | | | | | | | | |
| Resolution *(Å)* | 45.30-2.03 (2.08-2.03) | 48.79-1.79 (1.81-1.79) | 44.45-1.10 (1.11-1.10) | 45.84-1.30 (1.31-1.30) | 46.20-1.40 (1.42-1.40) | 47.61-1.35 (1.37-1.35) | 49.01-1.50 (1.52-1.50) | 45.88-1.35 (1.37-1.35) | 44.68-1.20 (1.21-1.20) | 44.56-1.35 (1.37-1.35) | 46.45-1.45 (1.47-1.45) |
| No. reflections | 38587 (3786) | 197115 (19666) | 221371 (20916) | 138104 (13378) | 111729 (11042) | 118571 (11677) | 90913 (8846) | 125082 (12359) | 177396 (17566) | 123977 (12210) | 101383 (10052) |
| $R_{work}/R_{free}$ | 0.198/0.240 (0.377/0.419) | 0.207/0.243 (0.363/0.392) | 0.161/0.173 (0.322/0.302) | 0.160/0.195 (0.343/0.381) | 0.178/0.203 (0.334/0.338) | 0.151/0.188 (0.299/0.339) | 0.158/0.182 (0.295/0.314) | 0.172/0.197 (0.308/0.290) | 0.146/0.167 (0.280/0.295) | 0.144/0.178 (0.269/0.296) | 0.165/0.194 (0.275/0.290) |
| No. atoms | 4692 | 18024 | 5539 | 5490 | 5345 | 5254 | 5303 | 5440 | 5602 | 5375 | 5445 |
| Protein | 4597 | 16915 | 4682 | 4774 | 4645 | 4650 | 4680 | 4765 | 4823 | 4746 | 4734 |
| Ligand/ion | 0 | 153 | 117 | 62 | 73 | 26 | 69 | 43 | 71 | 82 | 76 |
| Water | 95 | 956 | 740 | 654 | 627 | 578 | 554 | 632 | 708 | 547 | 635 |
| B-factors (overall) | 54.34 | 40.13 | 18.55 | 22.84 | 22.89 | 23.08 | 24.06 | 24.16 | 22.85 | 22.06 | 22.81 |
| Protein | 54.54 | 40.10 | 16.90 | 20.80 | 21.66 | 21.35 | 22.83 | 22.81 | 20.74 | 19.88 | 21.31 |
| Ligand/ion | - | 47.90 | 26.41 | 43.53 | 33.16 | 46.17 | 39.80 | 36.09 | 42.20 | 61.70 | 37.41 |
| Water | 44.77 | 39.44 | 27.76 | 35.80 | 30.75 | 35.91 | 32.51 | 33.55 | 35.30 | 35.05 | 32.20 |
| R.m.s. deviations | | | | | | | | | | | |
| Bond lengths *(Å)* | 0.007 | 0.006 | 0.005 | 0.006 | 0.006 | 0.006 | 0.006 | 0.006 | 0.005 | 0.006 | 0.005 |
| Bond angles *(°)* | 0.802 | 0.888 | 0.954 | 0.906 | 0.972 | 0.934 | 0.948 | 0.944 | 0.928 | 0.925 | 0.878 |

*Statistics for the highest resolution shells are shown in parentheses

**Supplementary table 2 –** Comparison of active site residues in orthologs. Cells are coloured by amino acid for ease of comparison. *or equivalent residue.

| **Residue number*** | *Lc* | *Cs* | *Ch* | *Md* | *Sc* | *Bd* | *Cc* | *Aa* | *Ag* | *Cq* |
|---|---|---|---|---|---|---|---|---|---|---|
| **1** | M | M | M | M | M | M | | | | |
| **3** | F | F | F | F | F | S | | | | |
| **100** | F | F | F | F | I | L | I | K | R | R |
| **137** | G/D | G | D | D | G | G | G | A | G | G |
| **139** | I | I | V | I | I | V | C | N | V | T |
| **140** | I | I | I | F | C | V | T | R | E | E |
| **147** | M | M | Y | W | Y | W | W | M | L | L |
| **148** | Y | Y | Y | F | Y | F | Y | Y | Y | Y |
| **308** | M | M | M | M | M | M | S | M | L | F |
| **309** | F | F | F | F | F | F | T | F | S | T |
| **354** | F | L | L | L | L | V | I | F | I | L |
| **355** | F | F | F | F | F | M | F | S | L | L |
| **420** | Y | Y | Y | Y | Y | L | Y | L | I | V |
| **457** | Y | Y | Y | Y | F | Y | Y | Y | Y | Y |
| **461** | R | R | R | R | R | R | R | F | M | M |

**Supplementary table 3:** Michaelis-Menten kinetic parameters for hydrolysis of diethyl umbelliferyl phosphate (organophosphate). Values are mean and standard error (n=4).

| Round of evolution | $k_5$ ($s^{-1}$) $\times 10^{-4}$ | $K_m$ (μM) | $k_5/K_m$ ($s^{-1}$ $M^{-1}$) |
|---|---|---|---|
| WT | 0.94±0.21 | <1.5 | >49 |
| R1 | 17±0.46 | <1.5 | >1100 |
| R2 | 43±0.59 | 1.9±1.3 | 2300±1600 |
| R3 | 61±0.45 | 6.4±0.97 | 960±150 |
| R4 | 58±2.5 | 3.0±0.75 | 2000±500 |
| R5 | 82±2.9 | 7.7±0.62 | 1100±91 |
| R6 | 69±0.76 | 6.5±0.56 | 1100±93 |
| R7 | 750±24 | 15±1.6 | 5100±590 |
| R8 | 640±19 | 9.1±0.60 | 7000±500 |
| R9 | 1040±56 | 49±11 | 2100±500 |

**Supplementary table 4:** Michaelis-Menten kinetic parameters for hydrolysis of 4-nitrophenyl butyrate (carboxylester). Values are mean ± standard error (n=4)

| Round of evolution | $k_{cat}$ ($s^{-1}$) | $K_m$ (μM) | $k_{cat}/K_m$ ($s^{-1}$ $M^{-1}$) $\times 10^4$ |
|---|---|---|---|
| WT | 52±1.5 | 46±3.4 | 1100±4.8 |
| R1 | 20±0.55 | 230±17 | 8.6±0.34 |
| R2 | 20±0.30 | 180±8.0 | 11±0.30 |
| R3 | 7.9±0.10 | 110±4.1 | 6.9±0.11 |
| R4 | 15±0.22 | 200±8.0 | 7.6±0.15 |
| R5 | 6.1±0.12 | 210±12 | 2.9±0.08 |
| R6 | 0.91±0.020 | 180±11 | 0.51±0.01 |
| R7 | 9.0±0.22 | 200±13 | 4.6±0.16 |
| R8 | 6.1±0.074 | 150±5.5 | 4.0±0.07 |
| R9 | 4.8±0.094 | 87±5.8 | 5.6±0.15 |

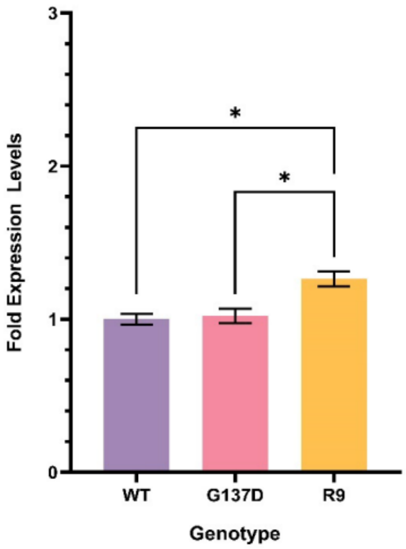


**Supplementary figure 3:** Relative expression levels of *Lcα*E7 transgenes. The expression of the *Lcα*E7 genes were normalised to the expression of housekeeping genes *Rpl11* and *CG13220*. The fold expression levels shown are relative to those measured for the WT gene. No expression of *Lcα*E7 was detected in the control line, hence the fold expression level cannot be calculated. The data represented: mean ± SEM; (n=3); Tukey's HSD test (P≤0.05 (*)).

**Supplementary table 5:** Toxicology data of Diazinon on *Drosophila melanogaster.* Sample size =50

| Diazinon | | | | | | | |
|---|---|---|---|---|---|---|---|
| Control | | WT | | G137D | | R9 | |
| Dose | Mortality | Dose | Mortality | Dose | Mortality | Dose | Mortality |
| 0 | 2 | 0 | 5 | 0 | 3 | 0 | 5 |
| 0 | 2 | 0 | 4 | 0 | 9 | 0 | 2 |
| 0 | 0 | 0 | 6 | 0 | 2 | 0 | 2 |
| 0 | 1 | 0 | 4 | 0 | 2 | 0 | 1 |
| 0 | 2 | 0 | 6 | 0 | 4 | 0 | 2 |
| 0.15 | 15 | 0.35 | 11 | 0.85 | 14 | 0.58 | 19 |
| 0.15 | 22 | 0.35 | 22 | 0.85 | 11 | 0.58 | 28 |
| 0.15 | 13 | 0.35 | 19 | 0.85 | 20 | 0.58 | 27 |
| 0.15 | 13 | 0.35 | 12 | 0.85 | 21 | 0.58 | 19 |
| 0.15 | 15 | 0.35 | 20 | 0.85 | 22 | 0.58 | 17 |
| 0.155 | 22 | 0.375 | 30 | 0.9 | 20 | 0.61 | 30 |
| 0.155 | 22 | 0.375 | 36 | 0.9 | 23 | 0.61 | 25 |
| 0.155 | 29 | 0.375 | 27 | 0.9 | 25 | 0.61 | 27 |
| 0.155 | 18 | 0.375 | 23 | 0.9 | 23 | 0.61 | 16 |
| 0.155 | 31 | 0.375 | 25 | 0.9 | 30 | 0.61 | 23 |
| 0.16 | 26 | 0.4 | 34 | 0.95 | 30 | 0.64 | 26 |
| 0.16 | 30 | 0.4 | 40 | 0.95 | 28 | 0.64 | 27 |
| 0.16 | 27 | 0.4 | 42 | 0.95 | 28 | 0.64 | 34 |
| 0.16 | 24 | 0.4 | 28 | 0.95 | 27 | 0.64 | 28 |
| 0.16 | 29 | 0.4 | 36 | 0.95 | 32 | 0.64 | 30 |
| 0.165 | 27 | 0.425 | 35 | 1 | 28 | 0.67 | 31 |
| 0.165 | 34 | 0.425 | 41 | 1 | 29 | 0.67 | 30 |
| 0.165 | 40 | 0.425 | 35 | 1 | 38 | 0.67 | 33 |
| 0.165 | 33 | 0.425 | 32 | 1 | 35 | 0.67 | 20 |
| 0.165 | 33 | 0.425 | 41 | 1 | 36 | 0.67 | 35 |
| 0.17 | 31 | 0.45 | 45 | 1.05 | 31 | 0.7 | 36 |
| 0.17 | 32 | 0.45 | 45 | 1.05 | 38 | 0.7 | 35 |
| 0.17 | 43 | 0.45 | 45 | 1.05 | 31 | 0.7 | 37 |
| 0.17 | 35 | 0.45 | 44 | 1.05 | 36 | 0.7 | 30 |
| 0.17 | 37 | 0.45 | 43 | 1.05 | 37 | 0.7 | 33 |

**Supplementary Table 6:** Toxicology data of Paraoxon on *Drosophila melanogaster.*
Sample size =50

| Paraoxon | | | | | | | |
|---|---|---|---|---|---|---|---|
| Control | | WT | | G137D | | R9 | |
| Dose | Mortality | Dose | Mortality | Dose | Mortality | Dose | Mortality |
| 0 | 1 | 0 | 4 | 0 | 5 | 0 | 5 |
| 0 | 3 | 0 | 5 | 0 | 1 | 0 | 5 |
| 0 | 3 | 0 | 3 | 0 | 3 | 0 | 2 |
| 0 | 1 | 0 | 2 | 0 | 3 | 0 | 5 |
| 0 | 4 | 0 | 6 | 0 | 3 | 0 | 1 |
| 0.058 | 8 | 0.112 | 15 | 0.55 | 17 | 0.5 | 17 |
| 0.058 | 15 | 0.112 | 13 | 0.55 | 12 | 0.5 | 18 |
| 0.058 | 13 | 0.112 | 22 | 0.55 | 11 | 0.5 | 13 |
| 0.058 | 20 | 0.112 | 27 | 0.55 | 16 | 0.5 | 9 |
| 0.058 | 10 | 0.112 | 12 | 0.55 | 18 | 0.5 | 15 |
| 0.062 | 17 | 0.118 | 27 | 0.6 | 19 | 0.55 | 19 |
| 0.062 | 17 | 0.118 | 31 | 0.6 | 22 | 0.55 | 17 |
| 0.062 | 22 | 0.118 | 31 | 0.6 | 21 | 0.55 | 27 |
| 0.062 | 21 | 0.118 | 23 | 0.6 | 26 | 0.55 | 22 |
| 0.062 | 12 | 0.118 | 22 | 0.6 | 19 | 0.55 | 22 |
| 0.066 | 19 | 0.124 | 36 | 0.65 | 23 | 0.6 | 25 |
| 0.066 | 30 | 0.124 | 40 | 0.65 | 24 | 0.6 | 29 |
| 0.066 | 28 | 0.124 | 41 | 0.65 | 19 | 0.6 | 26 |
| 0.066 | 25 | 0.124 | 38 | 0.65 | 23 | 0.6 | 27 |
| 0.066 | 20 | 0.124 | 30 | 0.65 | 28 | 0.6 | 35 |
| 0.07 | 31 | 0.13 | 42 | 0.7 | 32 | 0.65 | 39 |
| 0.07 | 35 | 0.13 | 40 | 0.7 | 33 | 0.65 | 34 |
| 0.07 | 38 | 0.13 | 45 | 0.7 | 35 | 0.65 | 36 |
| 0.07 | 27 | 0.13 | 47 | 0.7 | 32 | 0.65 | 31 |
| 0.07 | 21 | 0.13 | 33 | 0.7 | 31 | 0.65 | 36 |
| 0.074 | 34 | 0.136 | 42 | 0.75 | 46 | 0.7 | 42 |
| 0.074 | 37 | 0.136 | 45 | 0.75 | 35 | 0.7 | 43 |
| 0.074 | 42 | 0.136 | 43 | 0.75 | 39 | 0.7 | 40 |
| 0.074 | 17 | 0.136 | 48 | 0.75 | 40 | 0.7 | 42 |
| 0.074 | 34 | 0.136 | 35 | 0.75 | 35 | 0.7 | 40 |

**Supplementary table 7:** Primers and *LcαE7* sequences used

| Name | Sequence (5' to 3') |
|---|---|
| Real-Time Primers | |
| *LcαE7*_RT_F | ACCCACGCGAGATGGTTAAG |
| *LcαE7*_RT_R | TCTGCTTCAGGATGCTGGTG |
| *RPL11*_RT_F | CGATCCCTCCATCGGTATCT |
| *RPL11*_RT_R | AACCACTTCATGGCATCCTC |
| *CG13220*_RT_F | TGGGCAGTGCCTTCTACATTT |
| *CG13220*_RT_R | CGTACGCACCTCGCTTGTT |
| *Lcα*E7 WT & G137D Sequence (G137D mutation is underlined, GGC to GAT) | |
| 5' – ATGAACTTCAACGTGTCCCTGATGGAAAAGCTGAAGTGGAAGATCAAGTGTATCGAGAACAAGTTCCTGAACTACCGCCTGACC<br>ACCAACGAGACAGTGGTGGCCGAGACAGAGTACGGCAAAGTGAAGGGCGTGAAGCGCCTGACCGTGTACGATGATAGCTACTACAGCTTCG<br>AGGGCATCCCCTACGCTCAGCCACCAGTGGGAGAACTGCGTTTCAAGGCCCCACAGCGTCCAACACCATGGGATGGTGTTCGCGATTGCTGC<br>AACCACAAGGATAAGAGCGTGCAGGTCGACTTCATCACGGGCAAAGTGTGCGGCAGCGAGGATTGCCTGTACCTGAGCGTGTACACCAACAA<br>TCTGAACCCCGAGACAAAGCGCCCCGTGCTGGTGTACATTCACGGCGGA<u>GGC</u>TTCATCATCGGCGAGAACCACCGCGATATGTACGGCCCCG<br>ATTACTTCATCAAGAAAGACGTGGTGCTGATCAACATCCAGTACCGCCTGGGAGCCCTGGGATTCCTGAGTCTGAATAGCGAGGATCTGAACG<br>TGCCCGGCAATGCCGGCCTGAAGGATCAAGTGATGGCCCTGCGCTGGATCAAGAACAACTGCGCCAACTTCGGCGGCAACCCCGATAACATT<br>ACCGTGTTCGGAGAAAGCGCCGGTGCCGCCAGCACACACTATATGATGCTGACCGAGCAGACCCGCGGACTGTTTCACCGCGGTATTCTGAT<br>GAGCGGCAACGCGATCTGCCCCTGGGCTAATACTCAGTGCCAGCACCGCGCCTTCACACTGGCTAAGCTGGCTGGATACAAGGGCGAAGATA<br>ACGACAAGGACGTGCTCGAGTTCCTGATGAAGGCCAAGCCGCAGGATCTGATCAAGCTGGAAGAGAAGGTGCTGACCCTGGAAGAACGTAC<br>CAACAAAGTGATGTTCCCGTTCGGCCCCACCGTGGAACCATATCAGACAGCCGATTGCGTGCTGCCGAAGCACCCACGCGAGATGGTTAAGA<br>CAGCCTGGGGCAATAGCATCCCCACCATGATGGGCAACACCAGCTACGAGGGCCTGTTCTTCACCAGCATCCTGAAGCAGATGCCCATGCTG<br>GTCAAAGAACTGGAAACCTGCGTGAACTTCGTGCCCAGCGAACTGGCCGATGCCGAACGTACAGCCCCAGAGACACTGGAAATGGGCGCCA<br>AGATCAAGAAGGCCCACGTCACAGGCGAGACACCCACCGCCGACAACTTCATGGATCTGTGCAGCCACATCTACTTTTGGTTCCCCATGCACC<br>GCCTGCTGCAGCTGCGCTTCAATCACACAAGTGGCACCCCGGTGTACCTGTACCGCTTCGATTTCGATTCCGAGGACCTGATCAACCCCTACCG<br>CATCATGCGTAGTGGCCGCGGAGTGAAAGGTGTCAGCCATGCCGATGAGCTGACCTACTTCTTCTGGAACCAGCTGGCCAAGCGCATGCCCA<br>AAGAGAGCCGCGAGTACAAGACCATCGAGCGCATGACCGGCATCTGGATTCAGTTCGCCACCACCGGCAATCCCTACAGCAACGAGATCGAA<br>GGCATGGAAAACGTGTCCTGGGATCCGATCAAGAAAAGCGACGAGGTGTACAAGTGCCTCAACATCAGCGACGAGCTGAAGATGATCGATG<br>TGCCCGAGATGGACAAGATTAAGCAGTGGGAGTCCATGTTCGAAAAGCACCGCGATCTGTTCTAG – 3' | |
| *Lcα*E7 R9 Sequence | |
| 5' – ATGAACTTCAACGTGTCCCTGATGGAAAAGCTGAAGTGGAAGATCAAGTGTATCGAGAACAAGTTCCTGAACTACCGCCTGACC<br>ACCAACGAGACAGTGGTGGCCGAGACAGAGTACGGCAAAGTGAAGGGCGTGAAGCGCCTGACCGTGTACGATGATAGCTACTACAGCTTCG<br>AGGGCATCCCCTACGCTCAGCCACCAGTGGGAGAACTGCGTTTCAAGGCCCCACAGCGTCCAACACCATGGGATGGTGTTCGCGATTGCTGC<br>AACCACAAGGATAAGAGCGTGCAGGTCGACTTTATCGCCGGAAAAGTGTGCGGCAGCGAGGATTGCCTGTACCTGAGCGTGTACACCAACAA<br>TCTGAACCCCGAGACAAAGCGCCCCGTGCTGGTGTACATTCACGGCGGCGATTTCATCATCGGCGAGAACCACCGCGATATGTACGGCCCCG<br>ATTACTTCATCAAGAAAGACGTGGTGCTGATCAACATCCAGTACCGCCTGGGAGCCCTGGGATTCCTGAGTCTGAATAGCGAGGATCTGAACG<br>TGCCCGGCAATGCCGGCCTGAAGGATCAAGTGATGGCCCTGCGCTGGATCAAGAACAACTGCGCCAACTTCGGCGGCAACCCCGATAACATT<br>ACCGTGTTCGGAGAAAGCGCCGGTGCCGCCAGCACACACTATATGATGCTGACCGAGCAGACCCGCGGACTGTTTCACCGCGGTATTCTGAT<br>GAGCGGCAACGCGATCTGTCCATGGGCTAGTACACAGTGCCAGCACCGCGCCTTTACACTGGCCAAGCTGGCTGGATACAAGGGCGAAGATA<br>ACGACAAGGACGTGCTCGAGTTCCTGATGAAGGCCAAGCCGCAGGATCTGATCAAGCTGGAAGAGAAGGTGCTGACCCTGGAAGAACGTAC<br>CAATATGGTGGTGTTCCCGTTCGGCCCCACCGTGGAACCATATCAGACAGCCGATTGCGTGCTGCCGAAGCACCCACGCGAGATGGTTAAGA<br>CAGCCTGGGGCAATAGCATCCCCACCATGATGGGCAACACCAGCTACGAGGGCCTGTTCTTCACCAGCATCCTGAAGCAGACCCCGATGCTG<br>GTCAAAGAACTGGAAACCTGCGTGAACTTCGTGCCCAGCGAACTGGCCGATGCCGAACGTACAGCCCCAGAGACACTGGAAATGGGCGCCA<br>AGATCAAGAAGGCCCACGTCACAGGCGAGACACCCACCGCCGACAACTTCATGGATCTGTGCAGCCACATCTACTTTTGGTTCCCCATGCACC<br>GCCTGCTGCAGCTGCGCTTCAATCACACAAGTGGCACCCCGGTGTACCTGTACCGCTTCGATTTCGATTCCGAGGACCTGATCAACCCCTGGC<br>GCATTATGCGTAGTGGCCGCGGAGTGAAAGGCGTGGGACATGCCGATGAGCTGACCTACTTCTTCTGGAACCAGCTGGCGAAGCGCATGCCC<br>AAAGAGAGCCGCGAGTACAAGACCATCGAGCGCATGACCGGCATCTGGATTCAGTTCGCCACCACCGGCAATCCCTACAGCAACGAGATCGA<br>AGGCATGGAAAACGTGTCCTGGGATCCCGTGGAAAAGTCCGATGAGGTGTACCGCTGCCTGAACATCAGCGACGAGCTGAAGATGATCGAT<br>GTGCCCGAGATGGACAAGATTAAGCAGTGGGAGTCCATGTTCGAAAAGCACCGCGACCTGTTCTAA – 3' | |

**Supplementary table 8:** List of *Drosophila melanogaster lines*

| Lines | Description |
|---|---|
| 25709w | Contains 2nd Chromosome attP40 docking site for phiC31 integrase-mediated transformation (Bloomington Stock #25709) and an X-chromosome with a *white* mutation and phiC31 construct (Bloomington Stock #24749). |
| HR-GAL4 | Contains *Accord-GAL4* on 3rd Chromosome. |
| Transgenic Injected Lines (UAS- *LcαE7*) | |
| *UAS-Control* | Control line with *UAS-Null* on 2nd Chromosome. *UAS-Null* is a construct with no inserted gene. |
| *UAS-WT* | Line with *LcαE7 UAS-WT* on 2nd Chromosome. |
| *UAS-G137D* | Line with *LcαE7 UAS-G137D* on 2nd Chromosome. |
| *UAS-R9* | Line with *LcαE7 UAS-R9* on 2nd Chromosome. |
| Transgenic Overexpression Lines | |
| Control | Cross of *UAS-25709* with *HR-GAL4* lines. Control line with same genetic background as the overexpression lines. |
| WT | Cross of *UAS-WT* with *HR-GAL4* lines. *LcαE7* WT overexpression under the control of *Accord.* |
| G137D | Cross of *UAS-G137D* with *HR-GAL4* lines. *LcαE7* G137D overexpression under the control of *Accord.* |
| R9 | Cross of *UAS-R9* with *HR-GAL4* lines. *LcαE7* R9 overexpression under the control of *Accord.* |
